# A systematic comparison of VBM pipelines and their application to age prediction

**DOI:** 10.1101/2023.01.23.525151

**Authors:** Georgios Antonopoulos, Shammi More, Federico Raimondo, Simon B. Eickhoff, Felix Hoffstaedter, Kaustubh R. Patil

**Affiliations:** Institute of Systems Neuroscience, Heinrich Heine University Düsseldorf, Düsseldorf, Germany; Institute of Neuroscience and Medicine (INM-7: Brain and Behaviour), Research Centre Jülich, Jülich, Germany

## Abstract

Voxel-based morphometry (VBM) analysis is commonly used for localized quantification of gray matter volume (GMV). Several alternatives exist to implement a VBM pipeline. However, how these alternatives compare and their utility in applications, such as the estimation of aging effects, remain largely unclear. This leaves researchers wondering which VBM pipeline they should use for their project. In this study, we took a user-centric perspective and systematically compared five VBM pipelines, together with registration to either a general or a study-specific template, utilizing three large datasets (n>500 each). Considering the known effect of aging on GMV, we first compared the pipelines in their ability of individual-level age prediction and found markedly varied results. To examine whether these results arise from systematic differences between the pipelines, we classified them based on their GMVs, resulting in near-perfect accuracy. To gain deeper insights, we examined the impact of different VBM steps using the region-wise similarity between pipelines. The results revealed marked differences, largely driven by segmentation and registration steps. We observed large variability in subject-identification accuracies, highlighting the interpipeline differences in individual-level quantification of GMV. As a biologically meaningful criterion we correlated regional GMV with age. The results were in line with the age-prediction analysis, and two pipelines, CAT and the combination of fMRIPrep for tissue characterization with FSL for registration, reflected age information better.

## 1 Introduction

Analysis of brain structure has provided important insights regarding its organization in health and disease. T1-weighted (T1w) images obtained using magnetic resonance imaging (MRI) are commonly used for this purpose. However, raw T1w images cannot be compared directly due to their semiquantitative nature and inter- and intrasubject variability [1]. Volumetric analysis of T1w images using voxel-based morphometry (VBM) [2, 3] allows the investigation of the volumetric composition of brain tissues across subjects. It estimates tissue volume in each voxel and brings individual brains in a common reference space permitting comparison. VBM analysis has provided a plethora of valuable insights, for instance, in neurodegenerative diseases [4–8] and psychiatric disorders [9].

VBM has been successfully applied to study aging [10–12]. Recently, prediction of individuals’age based on VBM-derived information has proven to be a validated proxy for brain integrity and overall health [13–15], and promising for individualized clinical applications [14, 16–19]. Brain-age prediction is an important and widely studied topic that aims to estimate the trajectory of healthy brain aging [20, 21].

To estimate the GVM from T1w images, some specific steps must be performed. The main steps of a VBM pipeline are as follows: i) **Segmentation** creates probability maps where each voxel is assigned a probability of belonging to specific brain tissues, usually gray matter (GM), white matter (WM), and cerebrospinal fluid (CSF).**Brain extraction**, which is the process of removing the skull from an image and leaving only actual brain tissues and CSF, is also a segmentation process but in some cases is performed prior to segmentation of GM, WM and CSF.

**ii) Spatial registration/normalization** to a reference brain space is performed so that anatomical regions are aligned. The reference space can be either a general template (e.g., MNI-152) or a study-/data-specific template (henceforth referred to as data-template) [22– 24]. Data-templates are mainly used when comparing healthy subjects to patients to avoid bias due to general templates constructed from healthy populations. Several ways exist to create a data-template, and they are often created to match a standard space, such as the MNI space. Most VBM pipelines come with a general template.

**iii) Modulation** of the normalized tissue estimates aims at preserving the original amounts of tissue after spatial registration. To do so, normalized images are adjusted by the amount of local volume changes.

Since the introduction of VBM in 1995 [2], several alternatives and a multitude of options for each of the steps have been proposed. Even though various VBM pipelines utilize the same steps, the order of the steps may vary, and each step might use a different algorithm with several configurable options. Moreover, the pipelines can use those steps in a different order or perform some of them simultaneously and/or iteratively. It is also possible to create hybrid pipelines by combining the steps from different tools. Furthermore, optional steps, for example, whether to create a data template or use a general template provided by a tool, add to the already vast number of choices. Consequently, even if a user chooses an *off-the-shelf* VBM pipeline is not completely absolved of further choices. How the outputs of VBM pipelines compare and their utility in different applications remain poorly studied, which can lead to suboptimal choices [25–27].

Previous work comparing VBM pipelines indeed provides evidence for differences. A comprehensive comparison between Computational Anatomy Toolbox (CAT) [28] version 12.7, two FSL-based and a hybrid (still FSL [29] dependent) pipelines has shown that the choice of preprocessing pipeline has an impact both in age prediction and sex classification [30]. The same study showed that regions driving the results are pipeline dependent, while the choice of the templates used for registration, general or data-template, has little or no impact. FSL and SPM [31] yield different outcomes, especially for cortical regions [32]. A comparison focusing on registration and segmentation steps of SPM and FSL concluded that these preprocessing steps drive the regions identified in multiple amyotrophic lateral sclerosis [26]. Segmentation and registration as implemented in SPM8 newseg, SPM8 DARTEL [33], and FSLVBM were found to have substantial influence on GMV estimates and their relationship to age [34]. This study additionally concluded that pipelines with limited degrees of freedom for local deformations might overestimate between-group differences. Finally, the selection of tissue probability maps (TPMs) as priors for segmentation systematically impacts the segmentation outcome and, in turn, affects the statistical estimates [35]. The CAT12 VBM pipeline was found to perform better in the detection of volumetric alterations in temporal lobe epilepsy compared to the VBM8 toolbox [36, 37].

Several studies have investigated the effects of individual VBM steps and their parametrization. A comparison of 14 deformation algorithms used for registration found that SyN [38] from the Advance Normalization Toolkit (ANTs) [39] and DARTEL (CAT) were among those with the best performance, with SyN exhibiting the highest consistency across subjects [40] as well as being among the most robust to noise, partial volume effects and magnetic field inhomogeneities [41]. Segmentation algorithms from SPM, ANTs and FSL showed relatively small differences in controls, but significant differences appeared when comparing brains with atrophies, suggesting that the segmentation algorithm should be selected according to the brain characteristics of the study-population [42]. Dadar and colleagues compared six segmentation tools and confirmed significant differences between the tools as well as within-tool differences based on interscanner analysis [43]. For brain extraction, although FSL-BET has been reported to have low performance [42], it does not influence subsequent segmentation [44]. A comparison of SPM12, SPM8 and FreeSurfer5.3 [45] showed that SPM12 estimates of total intracranial volume (TIV) align better with manual segmentation [46]. SPM-based estimates in autism spectrum disorder and typically developing controls were closest to manual segmentation in terms of TIV, followed by FreeSurfer, while FSL appeared to underestimate TIV [47].

Taken together, different VBM pipelines produce different outcomes. The disagreement in VBM pipelines hinders precise localization and valid interpretation of tissue volume in the downstream analysis, e.g., atrophy in patients with multiple sclerosis [48–50]. To date, there is no standard method to calculate GMV or guidelines on which implementation of VBM is appropriate for a study at hand, e.g., age prediction. Additionally, the interaction of different algorithms and parameters in each step of VBM for estimating GMV and their effect on age estimates across the adult life-span, has not been thoroughly investigated. Moreover, the utility of a data-template created from healthy subjects and how it compares with a general template, especially in cross-site studies, remains unanswered. Here, to fill this gap, utilizing three large datasets (each n*>*500), we compared and evaluated five VBM pipelines including two *off-the-shelf* workflows and three modularly constructed pipelines utilizing commonly used neuroimaging tools. Each pipeline was implemented in two versions, one using a general template and one using a data-template, resulting in a total of 10 VBM pipelines. To remain consistent with our user-centric approach and developer guidelines, we adopted the default parameters unless there were specific recommendations from the developers [51]. First, we investigated whether different VBM pipelines produce GMV estimates that lead to different results in machine-learning-based predictions of individuals’ chronological age. We also calculated regional correlation to age, as GMV is known to decrease with age in healthy subjects. This extrinsic evaluation provides a more objective and utilitarian proxy for comparison [19, 20, 52, 53] and a criterion based on biological factors. Additionally, we showed that the pipelines indeed produce distinct patterns of GMV using machine-learningbased classification. Specifically, we address the following questions:

- How do the pipelines differ at the *region*- and the *subject-level*?
- What impact do *brain extraction,segmentation* and *registration* have on GMV?
- What is the effect of using a *data-template* compared to a *general template*?
- How do the pipeline outcomes compare in *univariate* and *multivariate* analyses?
- Which pipeline better reflects *brain aging* and performs best in *brain-age prediction*?

With this comprehensive and systematic comparative analysis of VBM pipelines, we aim to provide essential information and recommendations to researchers to help them select the VBM pipeline that best matches their research goals.

## 2 Materials and Methods

### 2.1 Datasets

We analyzed T1w images of healthy individuals from three large datasets covering the adult lifespan, eNKI [54]: population based sample of n=953 subjects, of which 573 had no psychiatric or neurological disorders or medication at the time of the scan (48.±17.2 years, 630 female). CamCAN [55, 56]: n=634 aging individuals without serious psychiatric conditions or cognitive impairment (54.8±=18.4 years, 320 female). IXI [57]: multisite sample of n=582 normal and healthy subjects (49.4±16.7 years, 324 female). (Table 5 in Supplementary Material)

### 2.2 Pipelines

CAT [28], a popularly used off-the-shelf VBM tool, is a successor of the first VBM pipeline implemented in SPM [3]. Here, we used the latest version CAT12.8 (r1813). Several general-purpose neuroimaging tools also provide functionality that can be used to create VBM pipelines. FSLVBM [58] uses tools from FSL [29] and is also widely used. ANTs [39] provides broad image processing and image analysis functionality, including all functions needed to perform VBM. Hybrid VBM pipelines that combine the functionality of different tools can be constructed, e.g., using fMRIPrep [59], which performs brain extraction using ANTs and then performs the rest of the steps using FSL.

We devised five VBM pipelines following the recommended steps and settings in the literature [39]: ANTs, ANTs-FSL, fMRIPrep-FSL, FSLVBM, and CAT. These pipelines were selected to reflect the choices that are common practice and easy to use. We used each pipeline with a standard template (the default templates for CAT and FSLVBM) irrespective of the dataset (general template) and with a dataset-specific template that was created and used for registration (data-template). Together, this resulted in ten pipelines.

#### 2.2.1 ANTs

We used ANTs version 2.2.0. First, each scan was corrected using the N4 bias field correction [60] and then segmented to select intracranial tissues using Atropos-based brain extraction [61]. Next, Atropos segmentation initialized with K-means was applied to segment the images into GM, WM and CSF. The GM-map images were registered to a template (general or data-specific) using a sequence of transformations. First, rigid body and affine transformations were applied, followed by a nonlinear BsplineSyN transform with the parameters set as in [62]. The Jacobian matrix from the spatial transformation was used to modulate the segmented GM. Data-specific templates were created using the ANTs build template method with default values. To create the template images, the transformations were averaged and used iteratively [39, 63]. To keep the template shape stable over multiple iterations of template building, the inverse average warp was calculated and applied to the template image.

To facilitate the analysis, the data-template process was initialized using a general MNI template. Therefore, the final data-template was also in the MNI space. For all processes requiring tissue masks and templates as well as for the registration to MNI, we used the ICBM 152 Non-linear Asymmetrical template version 2009a and corresponding tissue probability maps [64, 65].

#### 2.2.2 FSLVBM

We used FSL version 6.0. The images were prepared by automatically reorienting and then cropping part of the neck and lower head. Then, BET was used to extract the intracranial part of the brain, which was then segmented into GM, WM and CSF using FAST. Data-specific templates were created following FSLVBM’s process utilizing all GM images from a given dataset. GM segmented images were affinely registered to the ICBM-152 GM template, concatenated and averaged. This averaged image was then flipped along the x-axis, and the two mirror images were then reaveraged to obtain a first-pass, study-specific *af fine* GM template. Second, GM images were reregistered to this *af fine* GM template using nonlinear registration, averaged and flipped along the x-axis. Both mirror images were then averaged to create the final symmetric, study-specific,*non − linear* GM template. The resulting data-template was in the MNI space. The GM images were then nonlinearly registered to the template (either general or data-specific) and modulated. As the general template, we used the FSL-provided template (see Table 1).

**Table 1:**
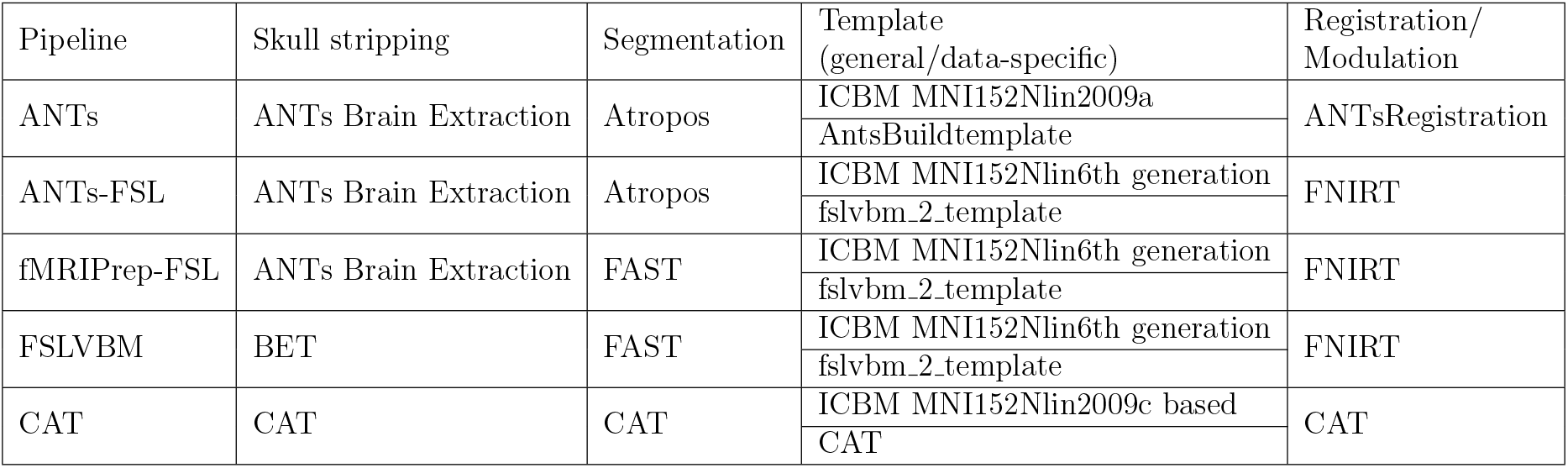
Software/algorithm used for the main VBM steps in our analysis pipelines.

#### 2.2.3 fMRIPrep-FSL

The reportedly poor quality of BET in brain extraction might lead to spurious results [42]; thus, we decided to test a pipeline that uses a better brain extraction as provided by ANTs followed by FSL for the rest of VBM processing. As fMRIPrep has been well validated and is gaining popularity, we chose to use the output of the fMRIPrep’s structural processing. In this hybrid pipeline for image preparation and segmentation, we used fMRIPrep version stable 20.0.6 [59], which uses ANTs version 2.1.0. Each T1w volume was corrected for intensity nonuniformity (INU) using N4BiasFieldCorrection [60] and skull-stripped using ‘antsBrainExtraction.sh‘ (using the OASIS template). Brain tissue segmentation into CSF, WM and GM was then performed using FSL FAST [66] (as used by the fMRIPrep FSL v5.0.9). This FAST parametrization diverges from the one in FSLVBM in the following parameters: (i) the Markov random field (MRF) beta value for the main segmentation phase was set to H=0.2, while the default value in FSLVBM was 0.1, and (ii) the MRF beta value for mixeltype was R=0.2, while the default in FSLVBM was 0.3. Template creation, spatial normalization, and modulation were identical to the FSLVBM pipeline.

#### 2.2.4 ANTs-FSL

The exact same processing, as mentioned above in the ANTs pipeline, was used to prepare the images, correct bias field noise, perform brain extraction and finally perform tissue segmentation using ANTs’ Atropos. The creation of a data-specific template, registration and modulation were implemented as in the FSLVBM pipeline. Note that the difference between this pipeline and the fMRIPrep-FSL pipeline is the tissue segmentation tool used.

#### 2.2.5 CAT

CAT12.8 was used based on SPM12 (v7771) using MATLAB (R2017b) and compiled for containerization in Singularity (2.6.1). CAT provides a complete VBM pipeline including denoising with spatial-adaptive nonlocal means, bias-correction, skull-stripping, and linear and nonlinear spatial registration. Images are segmented by an adaptive maximum a-posteriori approach [67] with partial volume model [68]. For nonlinear transformation, the geodesic shooting algorithm [69] is used. As the default template, an IXI-based template transformed to MNI152NLin2009cAsym is provided. For the data-template, initially, all structural T1 images are segmented into GM, WM, and CSF and spatially coregistered to the MNI standard template using affine registration. The affine tissue segments were used to create the new sample-specific geodesic shooting template that consists of four iterative nonlinear normalization steps.

Table 1 summarizes the VBM steps of each pipeline we utilized in our analyses.

#### 2.3 Parcellation scheme and quality control

To decrease the dimensionality of the data and thereby facilitate informative comparison and the use of machine-learning approaches, we extracted region-level averages. However, to preserve good spatial resolution, we selected a high granularity parcellation scheme. A combination of three atlases covering the whole brain and together constituting 1073 regions of interest (ROIs) was used: 1000 cortical regions from the Schaefer atlas [70], 36 subcortical regions from the Brainnetome Atlas [71] and 37 cerebellar regions [72]. Regional GMV values were calculated as the average of nonzero voxels within each region.

ANTs segmentation (Atropos), which was initiated with k-means, in some cases returned tissues in a different order, resulting in selecting the WM instead of the GM for further analysis. Therefore, we employed the following quality check to ensure that selected tissue represented GM. First, we discarded individuals who had a ratio of the mean of GM voxels over the mean of WM and CSF voxels of less than 1.5. Furthermore, images that were close to the 1.5 threshold as well as randomly sampled images were visually inspected for quality of segmentation. Because developing a thorough quality check or tackling this issue inside Atropos is out of the scope of this work, the threshold for the ratio of mean GM over WM and CSF was experimentally identified. Although CAT has an internal quality control method, for consistency, we applied our test to all pipelines. We retained only subjects who passed the quality checks across all the pipelines.

#### 2.4 Age prediction

We performed machine-learning-based analysis to predict the age of each subject using regional GMVs from each pipeline as features. We chose this as a suitable test given that age is reliably associated with GMV [19, 20, 52, 53] and because of the increasing importance of brain-age as a proxy for overall brain health [52, 73–75]. All features were standardized by removing the mean and scaling to unit variance in a cross-validation (CV)-consistent manner [76]. We utilized four machine-learning algorithms: relevance vector regression (RVR) [77], Gaussian process regression (GPR) [78], least absolute shrinkage and selection operator (LASSO) [79, 80], and kernel ridge regression (KRR) [81], in a nested 5-fold CV scheme repeated 5 times [82]. The age prediction performance was evaluated using the mean absolute error (MAE). To ensure that differences were not driven by factors other than the pipelines, we used the same data (subjects and regions) and models for each pipeline.

The evaluation was performed in two set ups, intradataset, and interdataset. In the interdataset evaluation, the models were trained using two datasets and then used to predict the third hold-out dataset. This analysis was performed for each pipeline separately.

#### 2.5 Classification of pipelines

To confirm the existence of systematic differences in the outcomes of the pipelines, we performed machine-learning-based predictive analysis based on the multivariate patterns of regional GMV. The idea behind this analysis is that if a model can classify the pipeline producing a GMV image with a high accuracy, that would indicate that the model learned systematic differences between the VBM pipelines. We performed 10-class classification with subjects’ regional GMVs as features and the pipelines as class labels. The features were standardized by removing the mean and scaling to unit variance in a CV-consistent manner [76] in two ways: i) within each feature and ii) within each subject. The former is standard preprocessing, while we implemented the latter to guard against trivial biases such as magnitude shifts. We used a linear support vector machine (SVM) with the default cost parameter of C=1 in a 5-fold CV scheme repeated 5 times.

#### 2.6 Individual-level identification

We examined the within-subject consistency of GMV patterns when processed by different pipelines. To do so, we identified subjects across pipelines using a nearest neighbor search. Using each pipeline as a reference (query), we tried to match each subject with all the subjects of each other pipeline (database). As an identification metric, we used Pearson’s correlation between two subjects’ regional GMVs [83, 84]. Each subject was matched with the subject from another pipeline with the highest correlation coefficient. The identification performance between two pipelines was calculated using the differential identifiability (Idiff) metric [84].

#### 2.7 Region-level comparison

To obtain a better understanding of regions driving the differences between pipelines, we assessed the similarity in regional GMV estimates from different pipelines using univariate statistical analysis. These analyses were performed for subjects from all datasets combined as well as separately for each dataset. We estimated similarity in regional GMVs across subjects using Pearson’s correlation coefficient for all possible pipeline pairs (in total 45). To investigate whether the size of parcels affects the regional similarities, we calculated for each ROI the median of correlation coefficients across the pairs of pipelines and correlated it with the number of voxels per region (see Figure 11 in the Supplementary Material).

For all arithmetic operations on Pearson’s *r* values,first Fisher’s *z* transform was applied, and then the result was transformed back to Pearson’s *r* value.

#### 2.8 Extrinsic evaluation of similarity between pipelines

The pipeline comparisons described above are intrinsic in nature. Thus, although they provide important information regarding differences between the pipelines, they do not provide information regarding the correctness of the pipelines in estimating the GMV. Such a correctness assessment, although desirable, cannot currently be achieved due to a lack of ground truth data. Instead, we compared the pipelines based on their utility in capturing age-related information.

We first tested to what degree regional GMV estimates from each pipeline reflect subjects’ age using univariate statistical analysis. To do so, we computed Pearson’s *r* between the regional GMVs and subjects’ ages for each pipeline separately. The resulting *p* values were corrected to control for the familywise error rate [85] due to multiple comparisons, again for all data combined as well as separately for each pipeline. We then performed an analysis of variance (ANOVA) to test whether the means of the correlation coefficients were significantly different.

Machine-learning-based analyses were performed using scikit-learn [86].

## 3 Results

### 3.1 Preprocessing and data-templates

For CAT and fMRIPrep, less than 0.4% of all subjects failed the preprocessing. For CAT, all outcomes passed our quality check. For FSLVBM, less than 2% of the subjects failed the QC. For fMRIPrep-FSL, there were slightly fewer subjects who failed QC than for FSLVBM. A considerable number of subjects failed ANTs segmentation (13% for eNKI, 5% for CamCAN and 12% for IXI). The QC results for the hybrid ANTs-FSL pipeline were similar to those of ANTs. The final number of subjects who qualified for further analyses was n=741 for eNKI, 593 for CamCAN and 418 for IXI (total n=1752).

The data-templates created by CAT and ANTs were sharper and more similar to general templates than those created by FSLVBM (templates are demonstrated in the Supplementary Material in Figures 8, 9, 10).

### 3.2 VBM pipelines produce different results

#### 3.2.1 Brain age prediction

We first performed individual-level prediction of chronological age using regional GMVs as features using four machine-learning algorithms (Figure 1). Within-dataset CV performance considerably varied among pipelines (Figure 1 (*a*)). The average performance across the learning algorithms and datasets was highest for the fMRIPrep-FSL general template (*MAE* = 5.83), followed by the FSLVBM general template (*MAE* = 6.17) and fMRIPrep-FSL data-template (*MAE* = 6.18). CAT with the data-template and with the general template showed similar performance of *MAE* = 6.37 and 6.39, respectively. The best average performance across datasets was achieved by the fMRIPrep-FSL general template with KRR (*MAE* = 5.59). ANTs performed the worst on average. All four learning algorithms generally showed similar performance for each pipeline (Supplementary Material Table 5).

**Figure 1:**
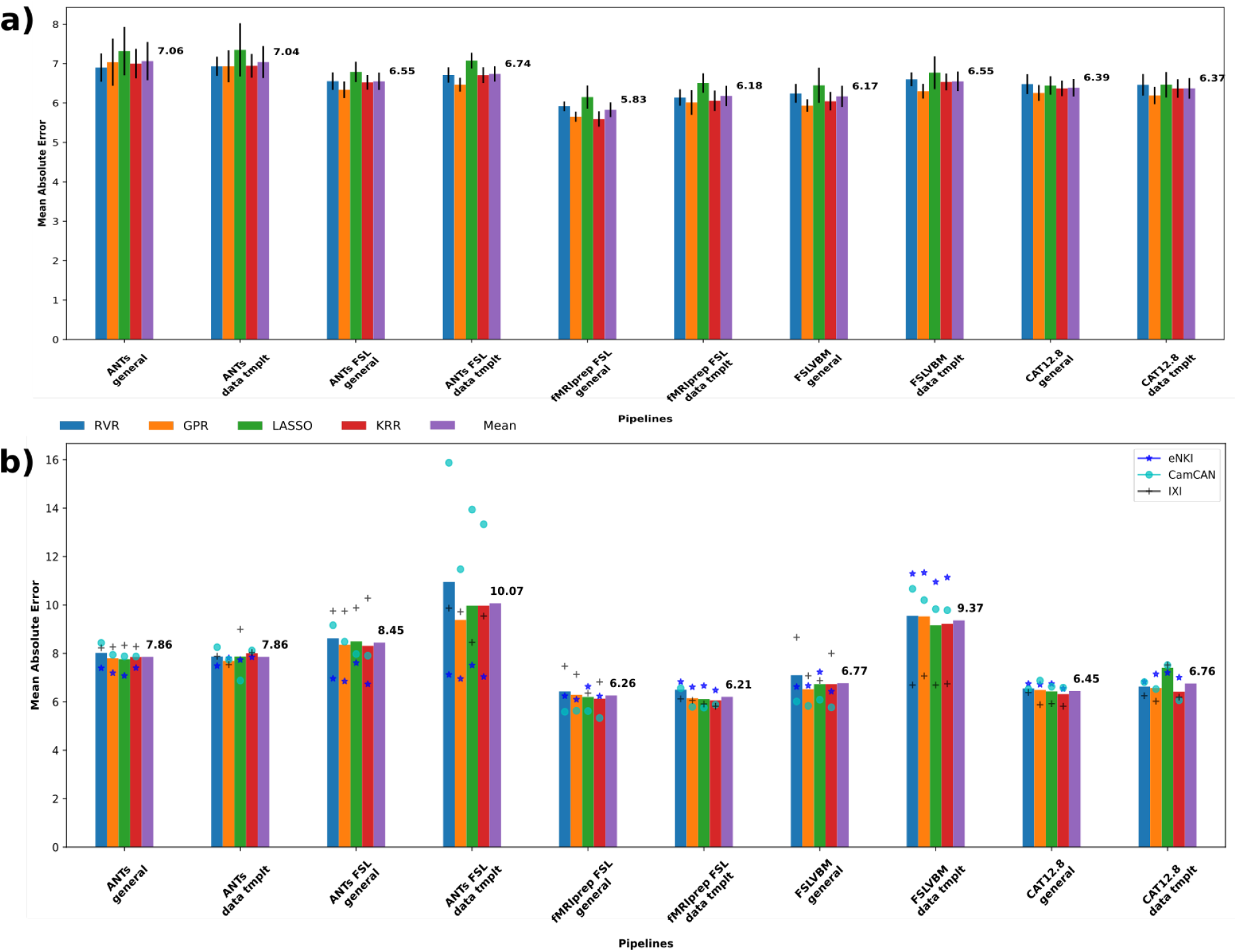
Age prediction for each pipeline. Blue, orange, green and red bars represent the averaged results of the three datasets per machine-learning algorithm, and the purple bars show the mean across models and datasets. a) Models trained and tested in the same dataset. Four models were tested using the three datasets in a nested K-fold cross-validation scheme. b) Age prediction for each pipeline when trained with two of the datasets and tested in the left-out one. Blue stars show the prediction performances on eNKI data, light blue circles the performances on CamCAN data, and black crosses on IXI data.

For cross-dataset predictions (Figure 1 (*b*)), the best performance averaged across datasets and models was again achieved by the fMRIPrep-FSL pipelines, with the data-template (*MAE* = 6.21) performing slightly better than the general template (*MAE* = 6.26) closely followed by CAT general template (*MAE* = 6.45). Here, the best overall predictions were again provided by the KRR algorithm. For the fMRIPrep-FSL data-template and generaltemplate *MAE* was 6.06 and 6.13, respectively. For CAT,*MAE* = 6.32 and 6.42 with the general template and data-template, respectively. ANTs-FSL-derived GMVs performed the worst on average (Supplementary Material Table 6).

#### 3.2.2 Machine-learning analysis confirms distinct GMV patterns

The machine-learning approach classified the pipelines with a near-perfect accuracy close to 100%. To rule out the possibility that this high accuracy was driven by systematic differences, that is, some pipelines over- or underestimating the GMV overall (which is indeed the case, see Supplementary Material Figure 12), we performed an additional analysis where each subject’s feature vector was *z*-scored independently, in effect removing the overall differences in GMV estimates. This analysis also resulted in high classification accuracy for all the datasets, close to 100%. Detailed results are provided in the Supplementary Material (Figure 13).

#### 3.2.3 Identification shows individual-level differences

Pipelines differing only in the template showed high differential identifiability 43*>*Idiff*>*29. fMRIPrep-FSL and FSLVBM, both with data-template, had the highest Idiff= 45, followed by the two ANTs pipelines (Idiff= 43). The two CAT pipelines had the lowest mean Idiff values, with the data-template pipeline being the lowest. FSLVBM with data-template had the highest mean Idiff. Pipelines using FSL for registration and modulation, with a general template, had a mean Idiff= 33.7. The same pipelines with a data-template showed mean Idiff= 37.7. ANTs-FSL and fMRIPrep-FSL, when both using a general template had Idiff= 35 and when using a data-template Idiff= 34. Finally, ANTs and ANTs-FSL, which differ in registration (and modulation), had Idiff= 29 when both used general templates and Idiff= 30 for data-templates (Figure 2).

**Figure 2:**
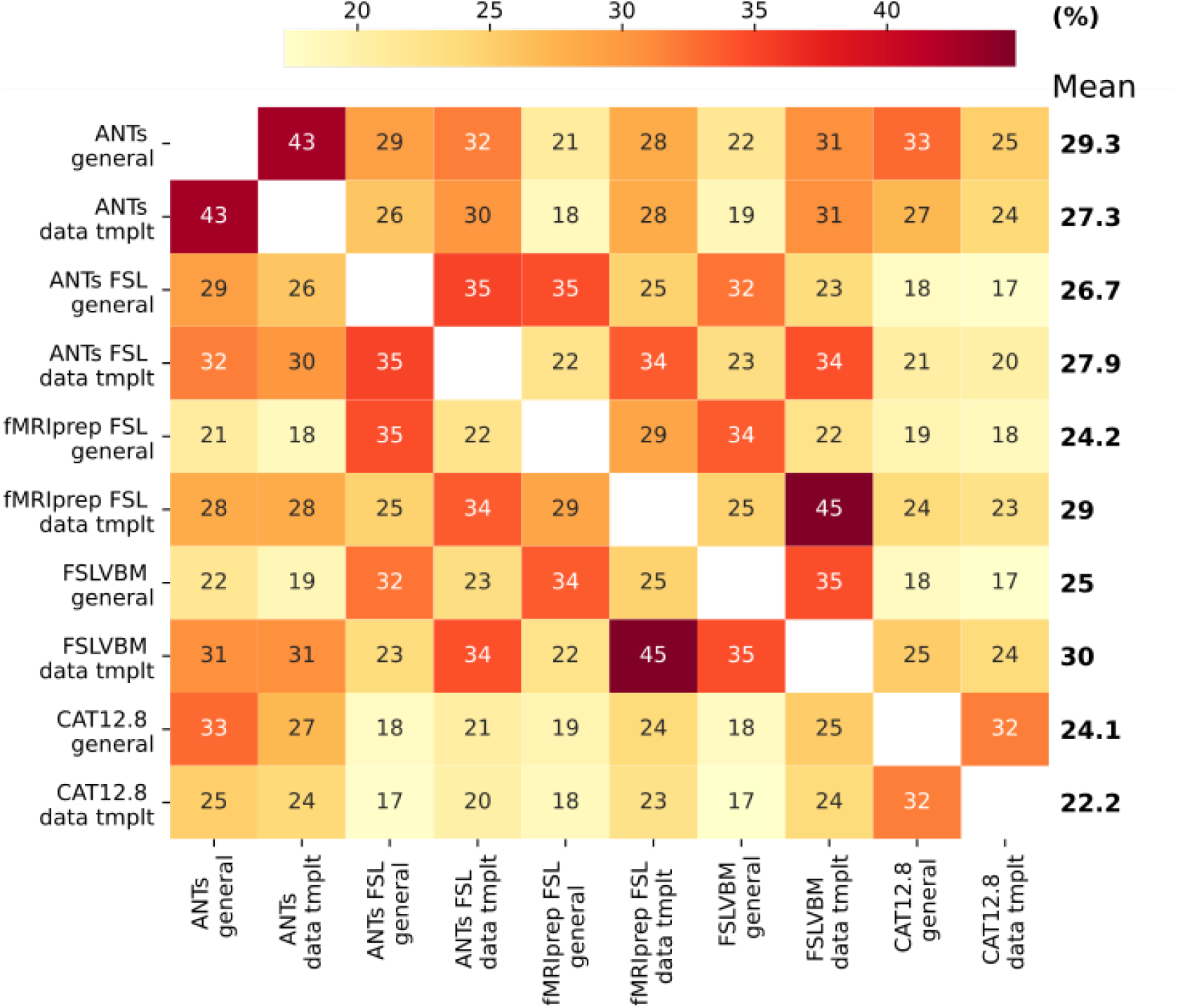
Identification performance in terms of differential identifiability. We used Pearson’s coefficient to calculate similarity between subjects. The highest mean Idiff was found for FSLVBM data-template followed by ANTs general template. The two CAT pipelines showed the lowest mean Idiff values.

#### 3.2.4 Univariate analysis and region-wise similarity

To better understand whether some VBM steps drive differences in the GMV estimates more than others, as well as to identify the regions showing significant differences, we performed several univariate statistical analyses. Some of the pipelines differ only in a single step; therefore, by examining the similarity between them, insightful conclusions can be extracted about the effect of this specific VBM step. We observed that the overall agreement between the pipelines, based on the median of the pairwise correlation values, varied across the regions, while most of the regions showed only low-to-moderate agreement (Figure 3). Only the regions close to the cingulum, temporal lobes and fusiform area showed relatively high agreement across the pipelines (median *r* >0.6). Most of the subcortical regions showed low agreement (median *r* < 0.4), except the caudate (median *r* >0.6). In the cerebellum, all regions showed a median *r* < 0.6. Overall, these results indicate a low agreement across the pipelines.

**Figure 3:**
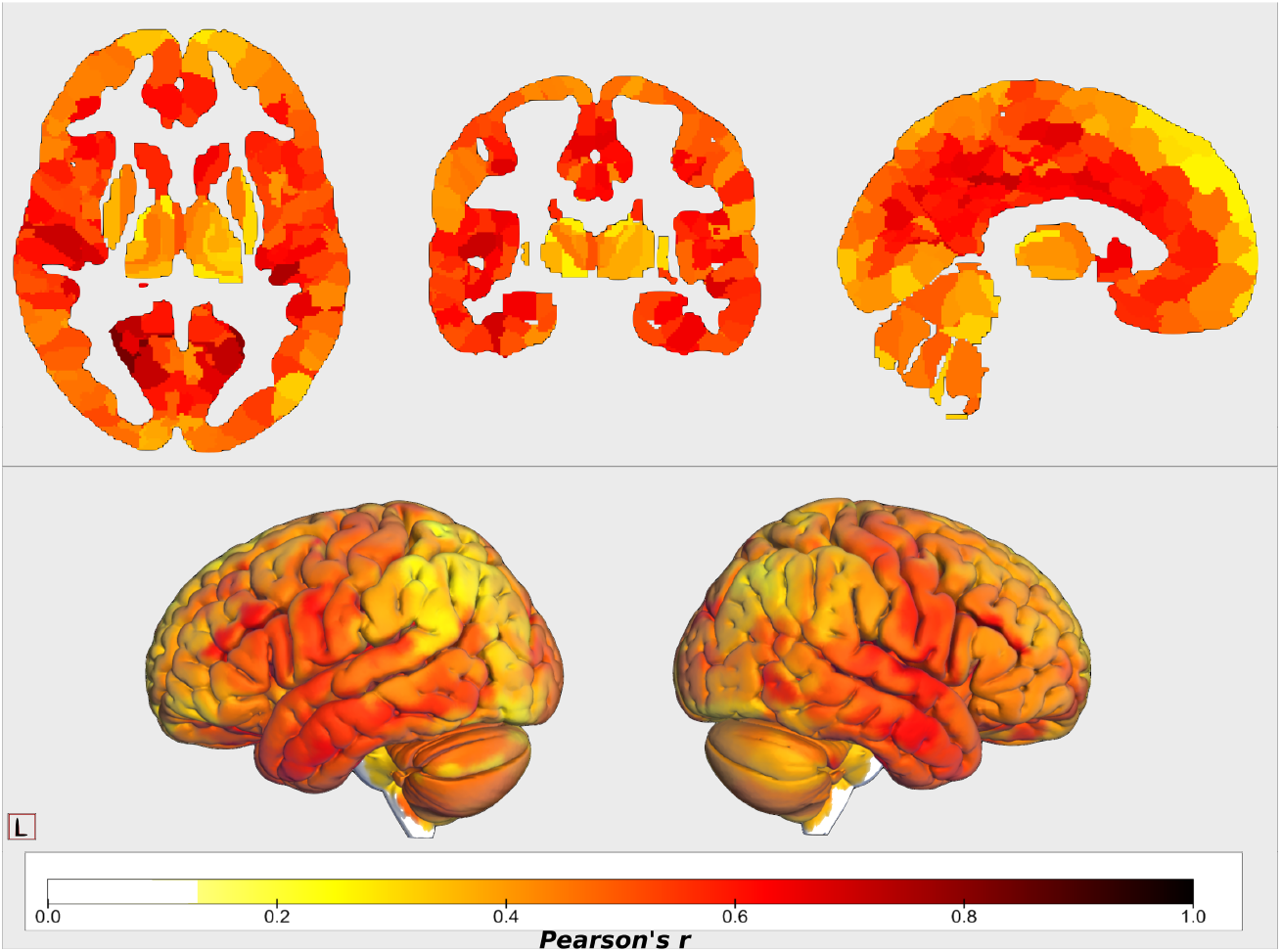
Median values of regional correlations calculated across subjects of all pairwise combinations of pipelines. The frontal lobe, subcortical regions and cerebellum showed lower similarity. First, correlations between regional GMVs across subjects were calculated for each pipeline pair. The median of these 45 values was then calculated as an overall agreement among the pipelines for each region.

The regionwise similarity between pairs of pipelines differed substantially. While ignoring pipeline pairs that differ only in the template (which are expected to be similar), maximum similarity was observed between fMRIPrep and FSLVBM both using a data-specific template (average *r* = 0.76), while the minimum similarity was between ANTs-FSL using the general template and CAT with both templates (average *r* = 0.306) (Figure 4).

**Figure 4:**
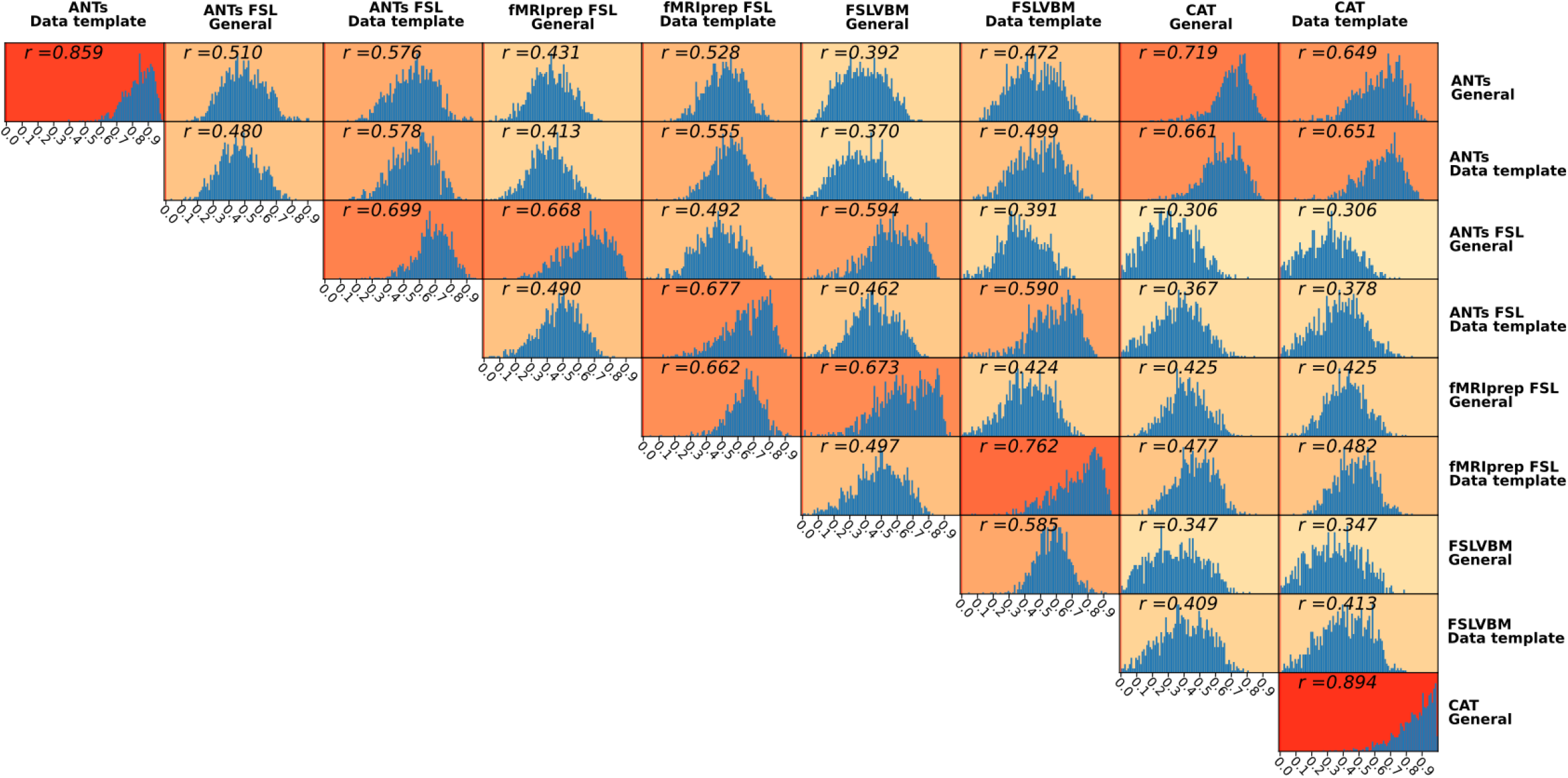
Histograms of regional interpipeline similarity for all pairs of pipelines. For each pair, we calculated Pearson’s r coefficient for each region across all subjects. We used the Holm-Bonferroni method to correct for multiple comparisons. The histograms shown consist of those regions that survived the multiple comparison (*p* < 0.05).

#### 3.2.5 Comparison between ANTs and CAT

High similarities were observed between the CAT and ANTs pipelines, despite differences in the steps, the order of the steps and the algorithms for each step. The highest similarity was observed when using the general templates (which themselves are different, as shown in Table 1) with *r* = 0.72 followed by *r* = 0.66 between the ANTs data-template and the CAT general template. A slightly lower similarity, of *r* = 0.65 was estimated when both pipelines used the data-templates as well as between the ANTs general template and the CAT data-template.

#### 3.2.6 Effect of Registration, Segmentation, and Brain extraction

In the subsequent analyses, we compared pipelines differing in specific VBM steps to assess their specific impact.

Regionwise similarity between ANTs and ANTs-FSL that differed only in **registration**(and therefore in modulation) using the general template was moderate to low, average *r* = 0.51. When using data-specific templates, the similarity was higher for all data (0.58) but also for each of the three datasets (Figure 5 (*a*)).

**Figure 5:**
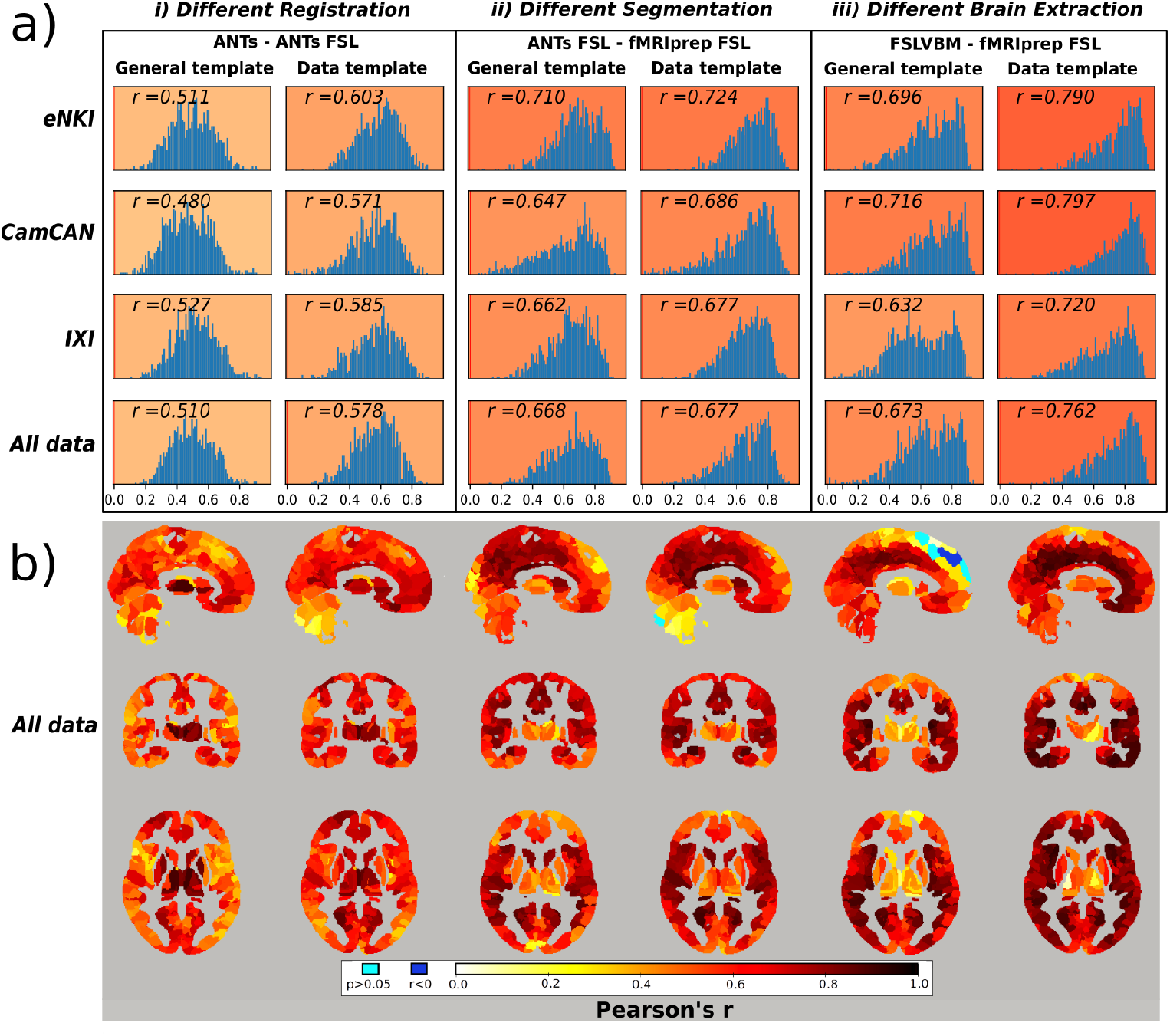
a) Histograms of regionwise correlation values between selected pairs of pipelines for all datasets. The *r* value represents the average correlation of all regions (that survived the Holm-Bonferroni correction) after transforming them to Fisher’s *z* and then reverse transformed to *r*. The pipeline pairs are categorized according to the template they use in the registration step. *i*) Correlation between ANTs and ANTs-FSL, which differ only in the registration step. *ii*) ANTs compared to fMRIPrep-FSL. These two pipelines differ only in the segmentation step, as fMRIPrep utilizes FSL-based segmentation. Segmentation imposes fewer differences than registration, *iii*) FSLVBM and fMRIPrep-FSL only differ in the brain extraction step. This step has a similar effect to segmentation when a general template is used and higher similarity when a data-template is used. The data-specific template comparisons are also provided here for convenience reasons, although it should be noted that the template creation steps may differ for the pipeline pairs, resulting in the usage of different data-specific templates. b) Brain maps with regional similarity of selected pairs of pipelines calculated using all data. Similarity values are expressed in Pearson’s r and were corrected using the Holm-Bonferroni method. Light blue represents regions without a significant association (p> 0.05) and blue represents regions with a negative correlation (*r* < 0).*i*) High similarity in subcortical areas and increased differences in cortical areas, especially when using a general template.*ii*) Different segmentations seem to have affected the cerebellum, subcortical areas and the posterior and anterior areas of the same axial level for both templates.*iii*) Brain extraction when using a general template caused more differences in the subcortical areas, superior frontal and the upper part of the cerebellum. It is noteworthy that negative values appear in the superior frontal lobe.

ANTs-FSL and fMRIPrep-FSL share the same steps besides **segmentation**. When using the general template, the average region-wise similarity was 0,67, and for the data-specific templates, the corresponding value was 0.68 (Figure 5 (*b*)).

FSLVBM and fMRIPrep-FSL differ in the **brain extraction** step. When both pipelines utilized the default FSL template, they had a similarity of 0.67. When the registration was performed using their respective data-specific template, the similarity increased to 0.76 (Figure 5 (*c*)).

Overall, similarities were higher when data-templates were used.

For ANTs compared to ANTs-FSL, the highest similarity values were in subcortical areas, and the lowest similarity values were in the ventrolateral and dorsolateral prefrontal cortices, especially when using a general template (Figure 5 *b*(*i*)). ANTs-FSL and fMRIPrep-FSL showed the least similarities in subcortical areas, the occipital lobe and prefrontal cortex (Figure 5 *b*(*ii*)). Finally, FSLVBM and fMRIPrep-FSL had the lowest similarity values in the subcortical areas, and the highest values were in the temporal lobes, medial prefrontal cortex and cingulate gyrus (Figure 5 *b*(*iii*)).

For each of the three datasets, similar figures separately with histograms of regional correlation values and Nifti files with all regional correlation values for the other pairs of pipelines can be found in the Supplementary Material.

#### 3.2.7 Pipelines with the same registration

ANTs-FSL and FSLVBM, which share only the registration step, had a similarity of 0.59 for all data when using either the FSL default or the data-specific template. The similarity for the eNKI dataset was 0.65 for both templates; for the CamCAN dataset, the similarity was 0.60 for the general template and 0.63 for the data-template and 0.56 and 0.58 for IXI dataset, respectively.

#### 3.2.8 General template versus data-specific template

The pipelines differing in the template, i.e., either general or a data-template, showed varying degrees of similarity (Table 2). The highest similarity was for CAT (*r* > 0.9), followed by ANTs (> 0.86) in all three datasets. The similarity was low to moderate for the three pipelines using FSL for registration and template creation steps (ANTs-FSL, FSLVBM, and fMRIPrep-FSL). Specifically, ANTs-FSL had a mean similarity across the three datasets of *r* = 0.71, fMRIPrep-FSL 0.66 and FSLVBM 0.59.

**Table 2:** The average values of regionwise correlation calculated across subjects for each pipeline when using a general template and a data-template. The *mean* across datasets is also presented, as well as the values from the same analysis performed with data from all datasets. It is noteworthy that when all data were combined, there was not an overall template created, but subjects were registered to the corresponding dataset template.

| General template compared to the data-specific template |  |  |  |  |  |
| --- | --- | --- | --- | --- | --- |
|  | ANTs | ANTs-FSL | fMRIPrep-FSL | FSLVBM | CAT |
| **eNKI** | 0.879 | 0.718 | 0.646 | 0.573 | 0.908 |
| **CamCAN** | 0.876 | 0.694 | 0.678 | 0.596 | 0.910 |
| **IXI** | 0.864 | 0.713 | 0.668 | 0.605 | 0.916 |
| **Mean** | 0.873 | 0.708 | 0.664 | 0.591 | 0.911 |
| **All data** | 0.859 | 0.699 | 0.662 | 0.585 | 0.894 |

Univariate analysis is in line with the identification Idiff results. Pearson’s r between the Idiff values and the regionwise correlations of pairs of pipelines was high, *r* = 0.841, *p* < 0.05 (more details in Supplementary Material Figure 18).

#### 3.3 Association with age

##### 3.3.1 Correlation between age and regional GMV

We performed univariate analysis to assess how regional GMVs capture aging-related information. CAT showed the highest average correlation magnitude between regional GMVs and age irrespective of the template used for all datasets, followed by fMRIPrep-FSL with the general template. For CAT, the mean correlation across datasets was *r* = − 0.410 and − 0.406 with a general template and data-specific template, respectively (Table 3). The distribution of regional GMV-age correlation values was more narrowly distributed for CAT and ANTs, while they were more broadly distributed for pipelines using FSL (Figure 6 (*a*)). Overall, the regional GMV-age correlation was markedly different between the pipelines (Figure 6).

**Table 3:**
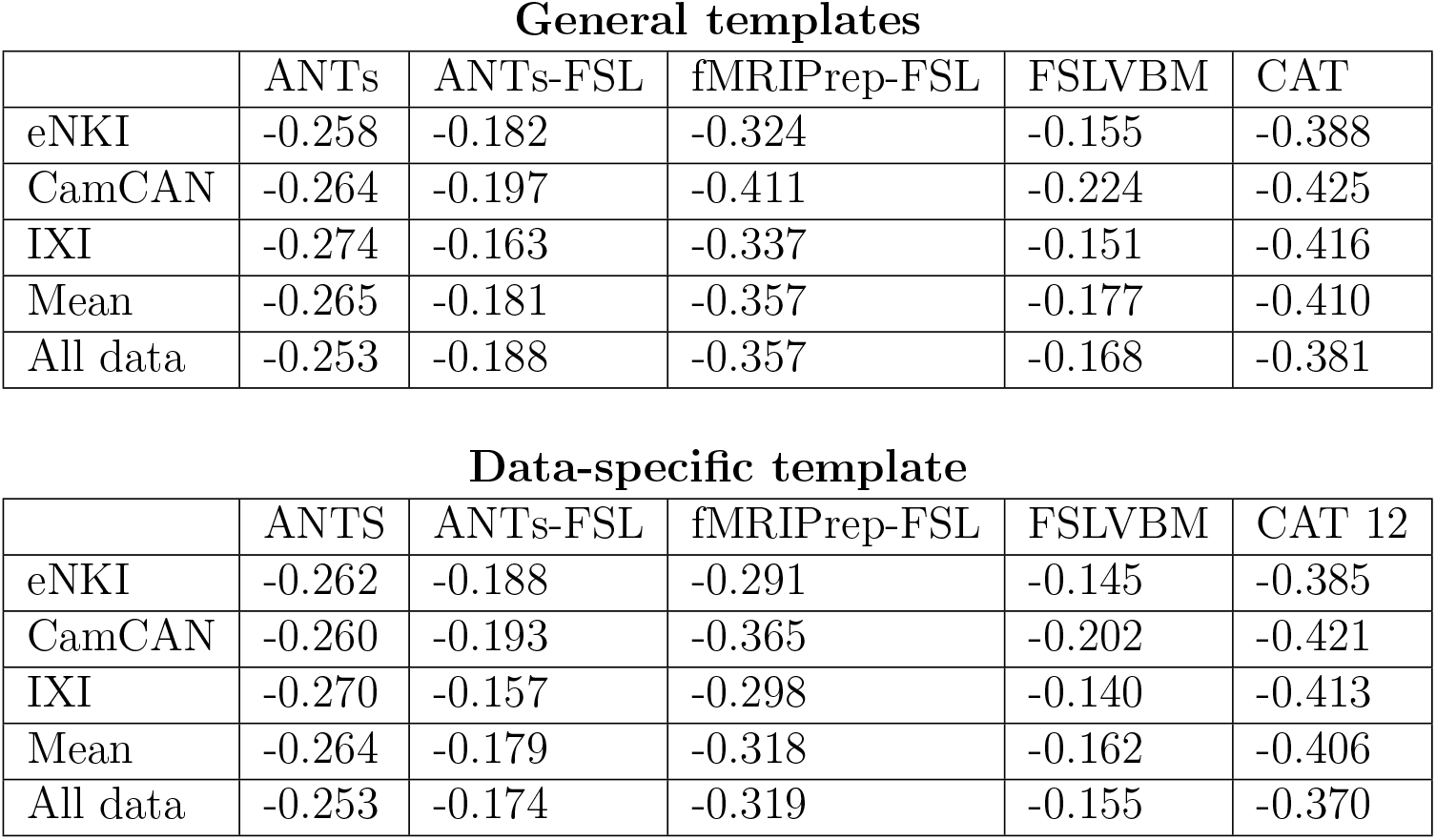
Pearson’s r-values were calculated between age and all regional GMVs across subjects. r-values were transformed to Fischer’s z averaged and transformed back to r-values. CAT with the general template and with the data-template appears to preserve age-related information better than the other pipelines, followed by fMRIPrep-FSL and ANTs. There is high consistency between datasets, with CamCAN showing a higher relation to age for those pipelines that use FSL for registration and CAT.

General templates
|  | ANTs | ANTs-FSL | fMRIPrep-FSL | FSLVBM | CAT |
| --- | --- | --- | --- | --- | --- |
| eNKI | -0.258 | -0.182 | -0.324 | -0.155 | -0.388 |
| CamCAN | -0.264 | -0.197 | -0.411 | -0.224 | -0.425 |
| IXI | -0.274 | -0.163 | -0.337 | -0.151 | -0.416 |
| Mean | -0.265 | -0.181 | -0.357 | -0.177 | -0.410 |
| All data | -0.253 | -0.188 | -0.357 | -0.168 | -0.381 |

Table 3: Pearson’s r-values were calculated between age and all regional GMVs across subjects.
|  | ANTS | ANTs-FSL | fMRIPrep-FSL | FSLVBM | CAT 12 |
| --- | --- | --- | --- | --- | --- |
| eNKI | -0.262 | -0.188 | -0.291 | -0.145 | -0.385 |
| CamCAN | -0.260 | -0.193 | -0.365 | -0.202 | -0.421 |
| IXI | -0.270 | -0.157 | -0.298 | -0.140 | -0.413 |
| Mean | -0.264 | -0.179 | -0.318 | -0.162 | -0.406 |
| All data | -0.253 | -0.174 | -0.319 | -0.155 | -0.370 |

**Figure 6:**
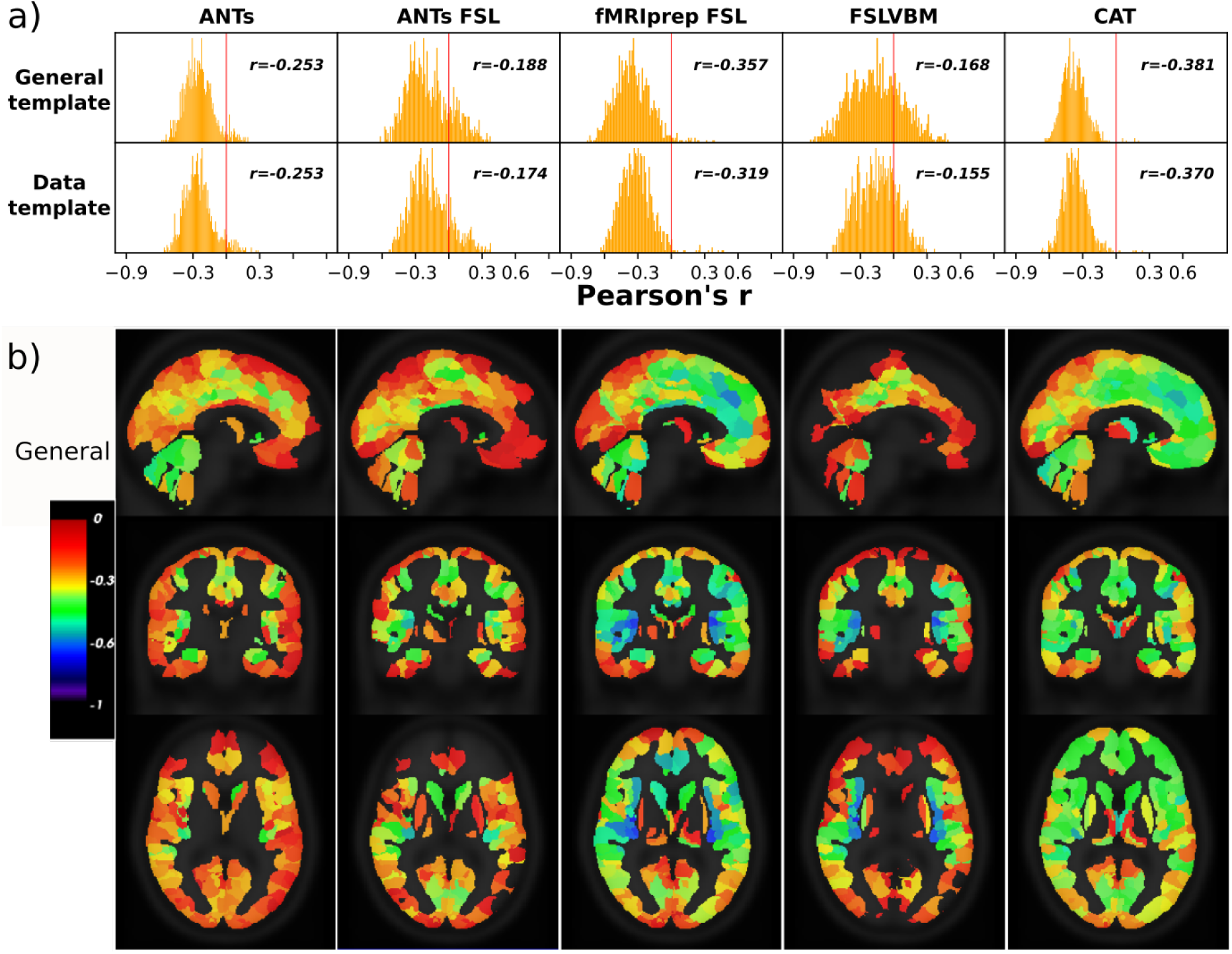
Correlation between regional GMV and age across subjects for the eNKI dataset. CAT had the fewest regions with a positive correlation with age (n=6 for the general template and 7 for the data-template). A few more regions with positive correlations had ANTs (n=27, n=31) and fMRIPrep-FSL (n=29 and 31). ANTs-FSL and FSLVBM have significantly higher numbers of regions with positive correlations as well as regions with non-significant correlations (*p*> 0.05). Regions with positive or nonsignificant correlations appear transparent in the brain images. For ANTs, the cerebellar regions and regions of cingulate gyri and limbic lobes. ANTs-FSL and FSLVBM demonstrated the most regions with a positive correlation with age. The cerebellum in FSLVBM shows a very small association with age, while in ANTs-FSL, cerebellar regions have more medium to high r values. Finally, fMRIPrep-FSL and CAT have small r values in the superior parietal and occipital lobes and medium to high r values in the frontal parts of the brain.

One-way ANOVA revealed a statistically significant difference in the average r-coefficients of regional GMV and age between at least two pipelines for all datasets (Supplementary Material Table 4).

##### 3.3.2 Comparison of regional age information between pipelines

The regional GMV-age correlation values not only differed but also showed opposing effects (Figure 7). In other words, some regions showed a positive correlation with age in one pipeline but a negative correlation in another pipeline (see Supplementary Material Figures 22, 23 and 24). In particular, this was the case for FSLVBM and ANTs-FSL, which contained many regions with a positive correlation with age. Strikingly, the same two pipelines also exhibited a large number of regions with opposing correlations with age when using a different template.

**Figure 7:**
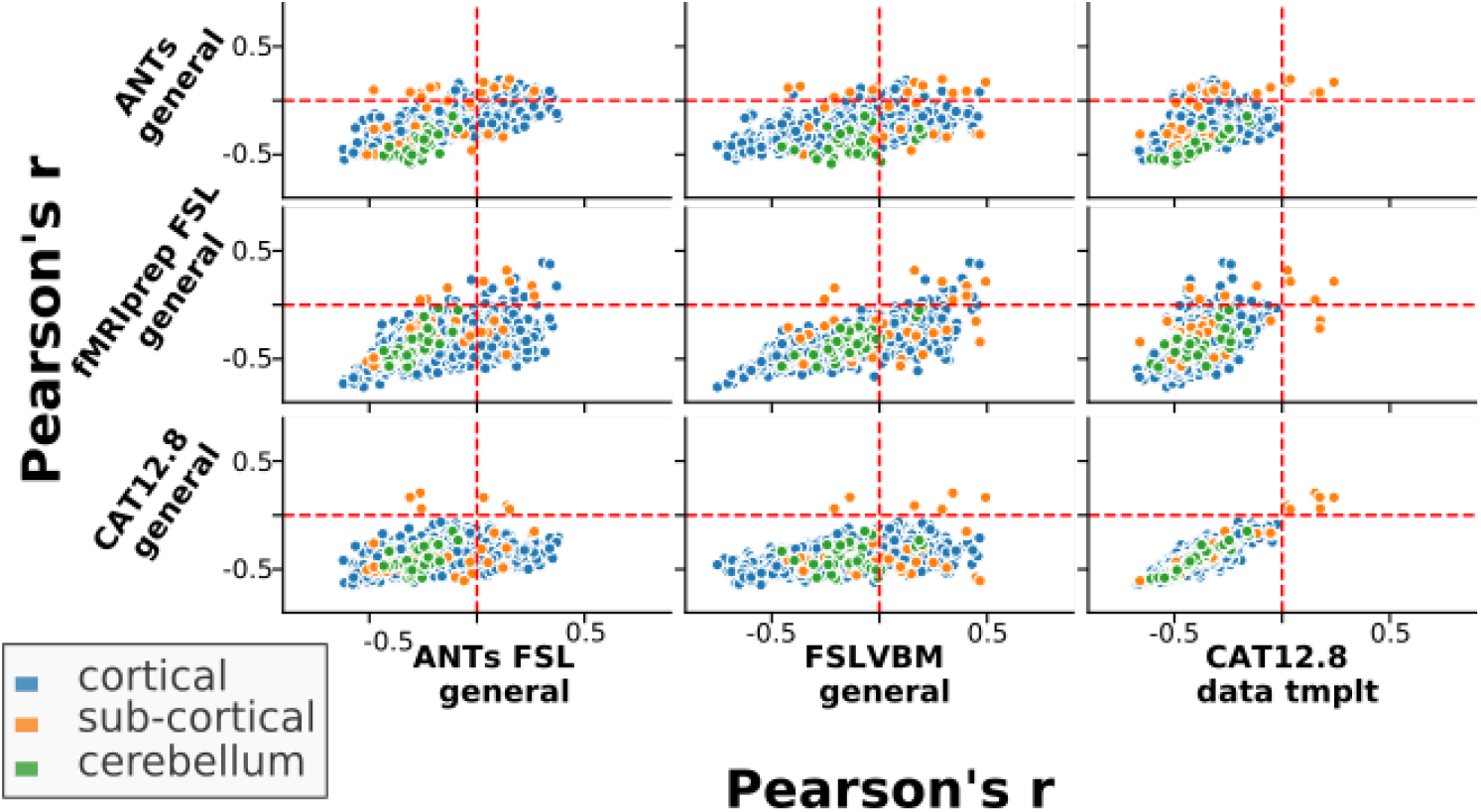
Pearson’s r values between regional GMV and age calculated across subjects for selected pipelines plotted against the same measurements for other pipelines. The upper left and lower right quadrants of each subplot contain those regions that have correlations to age with opposite signs/directions between the two pipelines. ANTs-FSL and FSLVBM have the most ROIs with positive correlations to age. Here, we selected a few pipelines that cover the spectrum of the main tools we used and better illustrate how the same regions in different pipelines can have opposite relations to age. All pipeline combinations can be seen in Figure 21 in the Supplementary Material.

When using all data, CAT had *n_rois* = 6 ROIs with a positive correlation to age when using either template. fMRIPrep-FSL had *n_rois* = 27 with the general template and 22 with the data-template, and ANTs had *n_rois* = 56 for both templates. ANTs-FSL and FSLVBM had *n_rois* = 218 and 280 regions positively correlated to age when using a general template and 184 and 226 regions when using a data-template, respectively. Two regions in the thalamus showed a positive correlation with age for all pipelines. In general, the regions with a positive correlation with age for all pipelines were mostly subcortical.

##### 3.3.3 Effect of Parcel size

We examined whether parcel size was associated with the agreement among the pipelines and with the agreement between ROIs and age. We observed no or marginal association between the overall similarity among the pipelines (calculated as the median of agreement between pipeline pairs) and parcel sizes (Pearson’s correlation, all data: *r* = − 0.08, *p* = 0.006, eNKI: *r* = − 0.02, *p* = 0.51, CamCAN: *r* = − 0.11, *p* = 0.0002, IXI: *r* = 0.07, *p* = 0.022)(Supplementary Material Figure 25).

Correlation values between parcel size and the corresponding regional correlation values to age for each pipeline varied between pipelines as well as between datasets. The highest correlation was for CAT, with *r* = −0.145 when using the general template and *r* = − 0.134 with the data-template (both *p* < 0.05). ANTs showed the next closest relation between parcel size and regional association with age, with *r* = − 0.105 when using a general template and *r* = − 0.101 when using a data-template (both *p* < 0.05). Those marginal negative correlations indicate that the fewer voxels are in an ROI, the better the relation of this ROI to age. All other correlation values were rather small, indicating that overall, the parcel sizes did not impact our results (Supplementary Material, for all data combined Figure 29, eNKI Figure 26, CamCAN Figure 27 and IXI Figure 28).

## Discussion

“Which tool shall I use to perform my VBM analysis?”, this is one of the very first questions that a researcher asks before starting a VBM study. The choice is often based on the literature or familiarity or recommendations. The current lack of an in depth comparison between VBM pipelines, the impact of the main steps on the outcome, and their utility precludes informative choice. Sparked by that, we compared 10 VBM pipelines derived from widely used tools on three large datasets covering the adult lifespan, acquired in different scanners and protocols. Two of the pipelines consisted of VBM steps from different tools. Our experiments were designed to facilitate a user-centric and systematic evaluation, which allows us to derive robust conclusions. Moreover, it permitted the examination of the effect of template use, i.e., general and data-template, as well as the effect of individual VBM steps.

Overall, we made the following observations based on analysis of the GMV estimates from different perspectives. The differences in individuals’ brain-age predictions confirmed that different VBM pipelines produce different GMVs (Figure 1, Tables 5 & 6). The systematic differences between the pipelines were further confirmed by the high accuracy when predicting the pipelines using their GMVs (Figure 13). A detailed univariate analysis of across-subject correlation (Figures 4) and identification using the subject-specific multivariate GMV pattern (Figures 2) showed that the individual steps of the VBM process as well as the choice of the template lead to the differences in the GMV estimates (see also Figure 5 and Table 2). Differences in GMV in turn impact the way age is reflected as we saw in univariate analysis correlating regional GMV with age (Figure 6 and Table 3).

First, we sought to establish whether the pipelines indeed lead to different results in applications. To this end, we performed predictive analysis using regional GMV as features and four machine-learning models commonly used in brain-age prediction. Individual-level age prediction showed variability in prediction accuracy (Figure 1), similar to what has been previously reported for voxel-level analysis and using CAT and FSL-based pipelines [30]. Our age-prediction accuracy for CAT and fMRIPrep-FSL are comparable to previous reports, considering our dataset size and the wide age range [87, 88]. To establish whether the differences in the pipelines are systematic, we performed classification analysis. The near-perfect classification performance in the prediction of pipelines (Figure 13) provides evidence for systematically distinct outcomes of the pipelines, which could be learned by the machine-learning algorithm and is in line with previous research [26, 32, 34]. Importantly, removing overall GMV differences by standardizing each feature vector also provided similarly high accuracy. Based on these results, even though the pipelines differ in seemingly trivial ways, such as using different templates or segmentation algorithm, we can conclude that they produce diverging GMV patterns.

The low to moderate identification performance and its variability across pipelines suggest that individual-level characteristics are, to a certain degree, captured differently by different pipelines (Figure 2). This result has important implications for data sharing and privacy issues [89]. As we show, with regionwise GMV data it is difficult to identify subjects when processed with different pipelines. Thus, when sharing such data, for instance, to perform multicenter analysis, it is important to keep the VBM pipeline consistent, including the template used.

Univariate analysis showed limited ROI-level similarity across pipelines, with an average regional similarity of *r* = 0.51 for pipelines using a general template. FSLVBM (using BET) and fMRIPrep-FSL (using ANTs brain extraction) showed high similarity, especially when a data-template was used (average *r* = 0.76) (Figure 5 (*c*)). When using the general template, the average similarity decreased but remained relatively high (*r* = 0.67). This suggests that differences in brain extraction are overshadowed by the subsequent steps. ANTs-FSL and fMRIPrep-FSL pipelines that differ mainly in segmentation (and the a priori template in brain extraction) showed relatively high agreement (*r* = 0.67 general template; *r* = 0.68 data-template), although slightly lower than what we show for brain extraction (Figure 5 (*b*)).

Differences between registration algorithms have been reported [41]. Our results are in line with this previous report. The registration step, evaluated as a comparison between ANTs and ANTs-FSL, had medium-to-high impact, with average agreement between these pipelines ranging across datasets, from *r* = 0.48 to *r* = 0.53 and *r* = 0.57 to *r* = 0.6 for general and data-template, respectively (Figure 5 (*a*)).

The impact of using different registration templates, general template versus data-template, was examined using pipelines that differ only in the template. This resulted in a wide-ranging agreement from *r* = 0.59 to *r* = 0.92 (Table 2). ANTs and CAT create data-templates that are very similar to their respective general templates – likely due to their exhaustive registration algorithms and the iterative processes together with the fact that their template creation processes are initialized with a general template. Overall, the differences in data-template creation algorithms and the ensuing data-templates led to substantial differences across the tools. This is in agreement with previous research reporting a small impact of the template when using CAT [35]. Effectively, using a data-template imposes higher similarity between the subjects’ images, which we also observed for some pipelines (Figure 4). Despite this high similarity, machine-learning-based analysis could reliably distinguish the pipelines. Univariate analysis of regionwise GMV-age correlations as well as age prediction were in favor of using a general template. Using subjects’ data to create a data-template and then registering the same subjects to it is a circular process unless an independent subset is used for template creation; however, given the limited data, this is often hard to implement in practice. The latter, in combination with the high computational demands of the template-creation process, are in favor of using a general template.

Although ANTs and CAT share no common modules, they showed medium to high similarity (for all data sets ranged from *r* = 0.65 to *r* = 0.72; maximum was for *r* = 0.74 for the eNKI). According to the impact of individual steps in the final GMV, as shown in our pipeline comparison, CAT and ANTs are expected to yield differing GMV estimates unless there are similarities in their internal algorithmic mechanism, which seems to be the case. In fact, exhaustive registration to similar templates can lead to similar outcomes. ANTs-FSL with the general template and CAT (both templates) showed the lowest regionwise similarity across datasets. However, in our opinion, the low similarity between CAT, with either template, and FSLVBM using a general template needs special attention (Figure 4 and Supplementary Material, eNKI Figure 14, CamCAN Figure 15 and IXI Figure 16). The reason is that they are both *off-the-shelf* pipelines and widely used in VBM projects. Regionally, the highest differences were present in the frontal lobe, superior parietal lobule and subcortical regions, specifically with regards to their association to age (Supplementary Material Figures 21, 22, 23, 24) Such differences enhance the risk of emanating different or even sometimes contradictory conclusions. From the projection of similarities between pipelines in the brain (Supplementary Material nifti files), it appears that high correlation values are not located in specific regions, nor is a specific pattern formed. However, segmentation and brain extraction seem to affect stronger subcortical and cerebellar areas and the superior frontal and occipital lobes. When comparing the registrations of ANTs and FNIRT, widespread differences appear in cortical areas and in the cerebellum (Figure 5 (*b*)).

The identification results (Figure 2) were very similar to the pairwise similarity estimated using Pearson’s correlation (Figure 4). The agreement between the two methods was high (Pearson’s correlation between pairwise similarity and Idiff, *r* = 0.84), and when using general templates, identification and univariate analysis were almost the same (*r* = 0.955, Supplementary Material Figure 18). This agreement between two different methods to assess similarity between the pipelines provides confirmatory validity to our findings.

It is important to note that, mostly for brain extraction but also for segmentation and registration algorithms, there are important differences between the datasets (Figure 5). This indicates that properties such as the intensity range of the images can influence the results in different ways, e.g., the quality of segmentation varies across different scanning parameters [90–92].

By using three large datasets, we aimed to cover a wide range of MRI vendors as well as scanning parameters and settings. Different scanners were used not only across datasets but also within the same dataset, strengthening our results and conclusions independent of the datasets’ idiosyncrasies.

The fMRIPrep-FSL combination showed the second highest correlation with age and the best brain-age predictions. This is not surprising given the nonexhaustive registration of FSL, which together with deep neural networks provides accurate brain-age prediction [25]. It is noteworthy that we used all subjects from the eNKI sample without separating the healthy part of the cohort as is usually done. When inspecting the age predictions of only healthy subjects, in intrasite predictions, and a mix of healthy and nonhealthy subjects, cross-site, separately, we did not observe a significant difference (see Supplementary Material Table 5 and Table 6). This can be explained by the fact that the nonlinear transformations wipe-out small differences compared to linear registration but also by the fact that the templates we used are based on healthy populations. In the age-prediction CAT showed performance similar to fMRIPrep-FSL but lower than what has been previously reported [17]. However, this difference can be driven by the machine-learning algorithms and the feature space employed. These results are in line with the univariate analysis we performed, where the same two pipelines had the highest (anti-) correlation with age (Figure 6). In addition, fewer ROIs showed a positive correlation with age for CAT and fMRIPrep-FSL than for other pipelines, which is in line with known GM atrophy with age [93–95]. Taken together, our results are in favor of CAT and fMRIPrep-FSL in regard to aging-related studies. Although some recent brain-age applications have shown that linear registration is preferable [16, 25], we decided to compare the whole VBM process using nonlinear registration. This choice was made so that we could approach the topic via a common space, permit the use of a parcellation atlas and facilitate the interpretability of the results.

The user-centric approach we followed in this project does not allow for an extensive evaluation of the potentials of the tools we used. CAT, ANTs, but to a certain degree also FSLVBM potentially can be tuned to provide more accurate brain-age predictions or regional associations to age. However, such an investigation is out of the scope of this work.

To summarize, our results show that all steps of a VBM pipeline have a considerable impact on the GMV estimates, and therefore, different pipelines produce different results. These differences in GMV estimates are reflected in univariate as well as multivariate analyses. The choice of registration has the highest impact, followed by segmentation and brain extraction algorithm. In the specific case of age-prediction, we recommend the combination of ANTs for brain extraction and FSL for segmentation (as implemented in fMRIPrep) and FSL nonlinear registration or CAT 12.8, with the latter having the advantage of being available as an off-the-self pipeline. The option of using a general template is preferred for age-related studies and likely other studies with a similar set up, especially when analyzing scans from multiple datasets.

## Ethics statement

Ethical approval and informed consent were obtained locally for each study covering both participation and subsequent data sharing. The ethics proposals for the use and retrospective analyses of the datasets were approved by the Ethics Committee of the Medical Faculty at the Heinrich-Heine-University Düsseldorf.

## Data/code availability statement

The codes used for preprocessing, feature extraction and model training are available at https://github.com/juaml/VBM_comparison.

## 5 Acknowledgments

This study is supported by Deutsche Forschungsgemeinschaft (DFG, PA 3634/1-1 and EI 816/21-1), the National Institute of Mental Health, the Helmholtz Portfolio Theme “Super-computing and Modelling for the Human Brain” and the European Union’s Horizon 2020 Research and Innovation Program grant agreement 945539 (HBP SGA3)

