## Supplementary Material for "A systematic comparison of VBM pipelines and their application to age prediction"

### Appendix

- MRI acquisition details

| Sample | Scanner | Sequence | Tesla | Slices | Voxel size (mm) | Time parameters (TR / TE / TI [ms]) | Other parameters (FA/FOV [ °/mm]) |
|---|---|---|---|---|---|---|---|
| eNKI | Siemens Magnetom TrioTim | 3D MP-RAGE | 3T | 176 | 1 x 1 x 1 | 1900/2.52/900 | 9/250x250 |
| Cam-CAN | Tim Trio Siemens | 3D MP-RAGE | 3.0T | 192 | 1 x 1 x 1 | 2,250/2.99/900 | 9/256 x 256 |
| IXI-Guys | Phillips | 3D MP-RAGE | 1.5T | 192 | 1.2 x 0.94 x 0.94 | 9.813/4.603/ | 8/256 x 256 |
| IXI-HH | Discovery GE | 3D FSPGR | 3T | 176 | 1.2 x 0.94 x 0.94 | 9.6/4.6/450 | N/A /256 x 256 |
| IXI-IOP | GE | N/A | 1.5 | N/A | 1.2 x 0.94 x 0.94 | N/A | N/A /256 x 256 |

- Preprocessing

All pipelines were run in a high-throughput compute cluster except CAT12.8, which was run in a high-performance computing cluster.

- Study specific templates of each pipeline

The templates created by CAT and ANTs appear to be sharper than those created by FSLVBM. For ANTs, the template was 197x233x189; for CAT 175x199x175; and for FSL-based pipelines, 91x109x91. Templates built with eNKI are presented in Figure 8, and templates built based on CamCAN subjects can be found in 9 and for IXI in 10 .

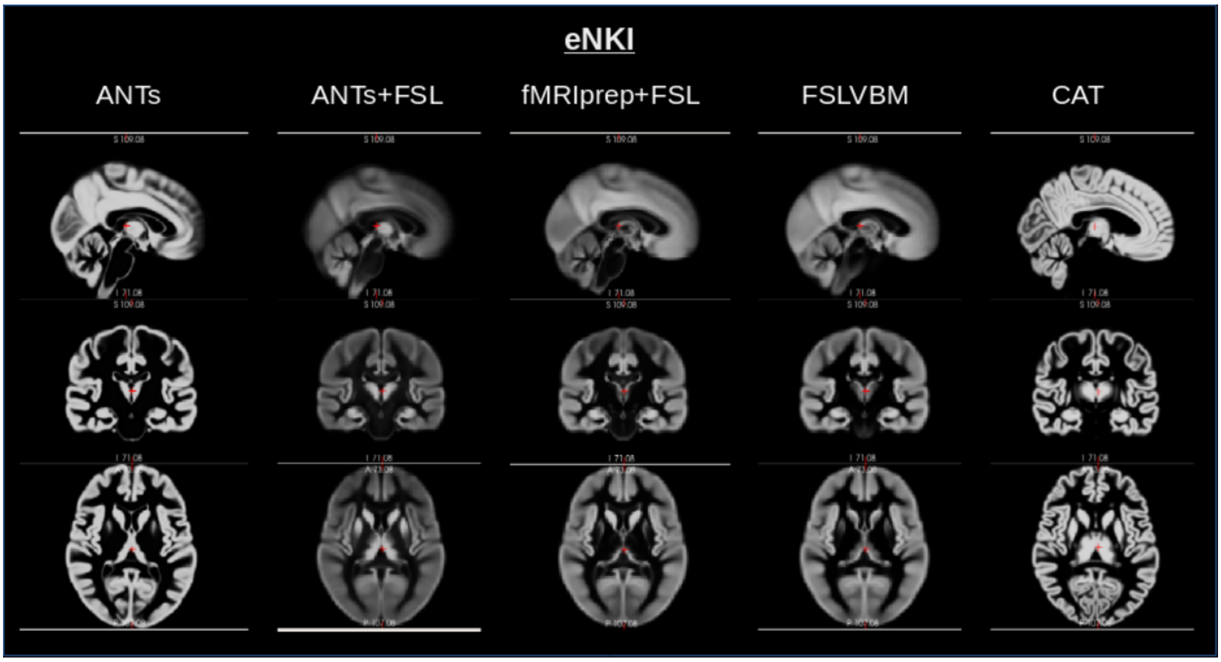


Figure 8: Templates created for each pipeline. Default CAT and ANTs processes created sharper templates.

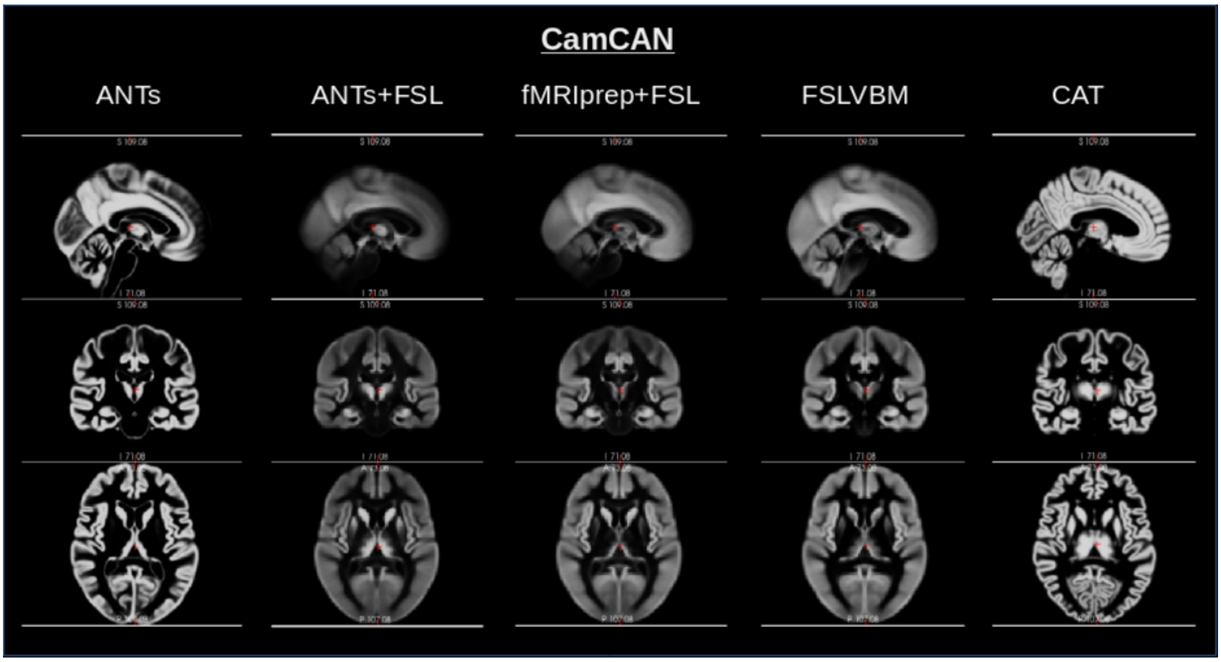


Figure 9: Templates created for each pipeline CamCAN dataset.

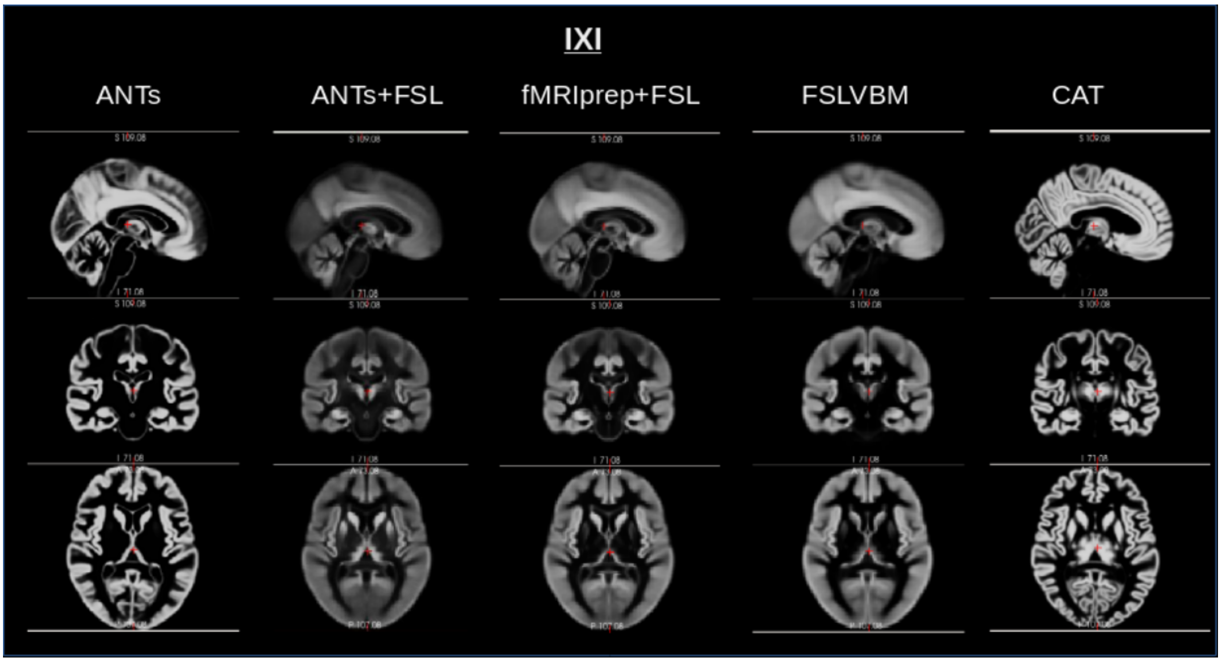


Figure 10: Templates created for each pipeline for the IXI dataset

- Initialization of ANTs Atropos

It is worth mentioning that to follow our user-centric perspective, we used K-means clustering to initialize ANTs Atropos segmentation instead of some a-priori probability map. This resulted in relatively many subjects failing the preprocessing, as tissue samples were assigned unexpected labels by the K-means algorithm, resulting in confusion between white matter and gray matter. Although this could be approached by adjusting Atropos, such effort is out of scope of this project; instead, we performed custom quality control to detect such failed preprocessing.

- Univariate analysis chart

Figure 11 illustrates the univariate analysis we followed.

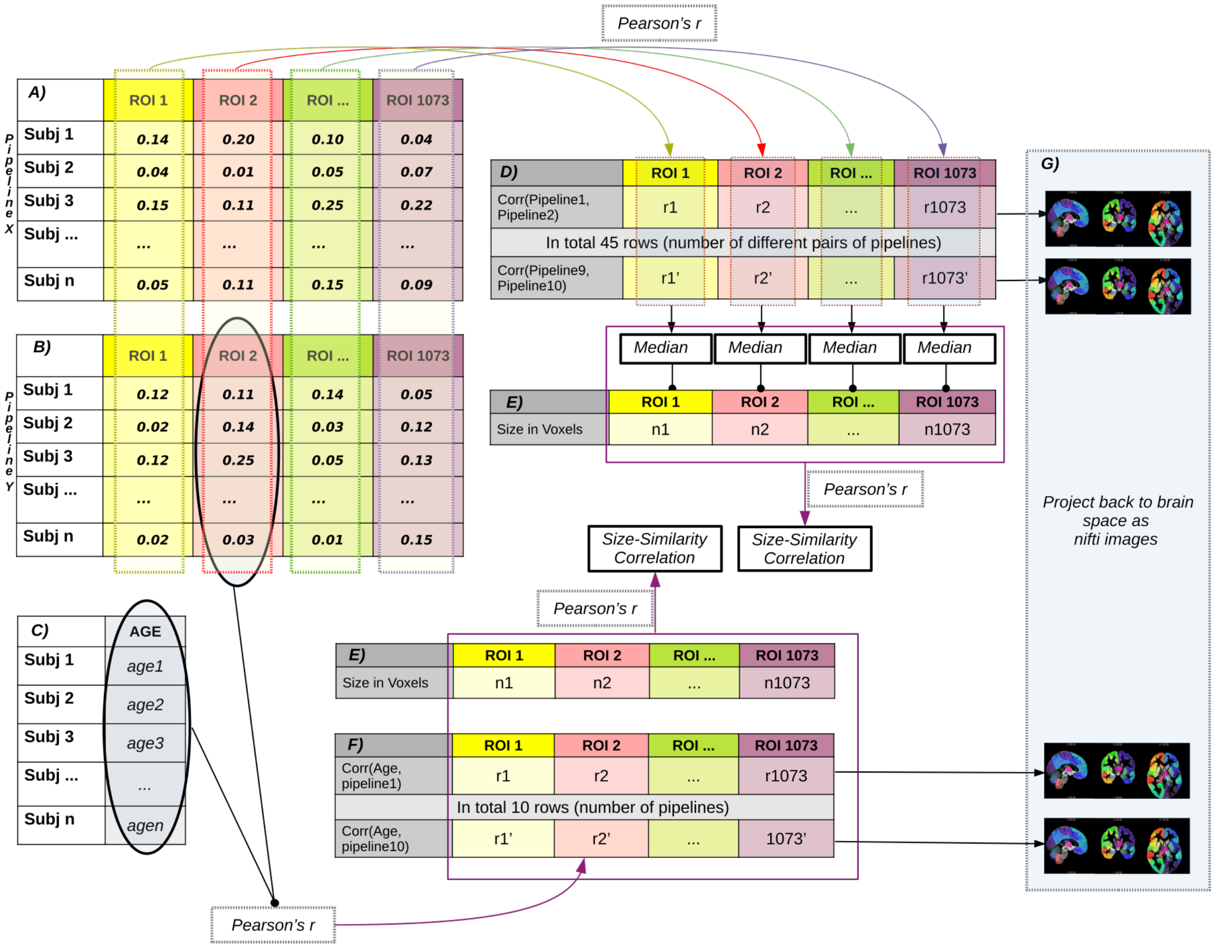


Figure 11: Depiction of the process to calculate regional correlations across subjects for a pipeline pair. For a given pair of pipelines (panels $A$ and $B$ in the figure) for each region, we calculate the correlation across subjects. By performing this process for all pairs of pipelines and all regions, we obtain a regional correlation matrix (panel $D$). The overall agreement between the pipelines was calculated as the median for each region across all pipeline pairs, which was then used to correlate with the size of the parcels (panel $E$). For each pipeline and each region, we calculated Pearson's correlation across subjects between regional GMV (shown here for panel $B$) and age (panel $C$). Regional correlation values between pipelines (panel $D$) or with age (panel $F$) were projected on the brain for visualization purposes.

#### Total GMV plots

The following image 12 presents the total GMV of all subjects for each pipeline and each dataset.

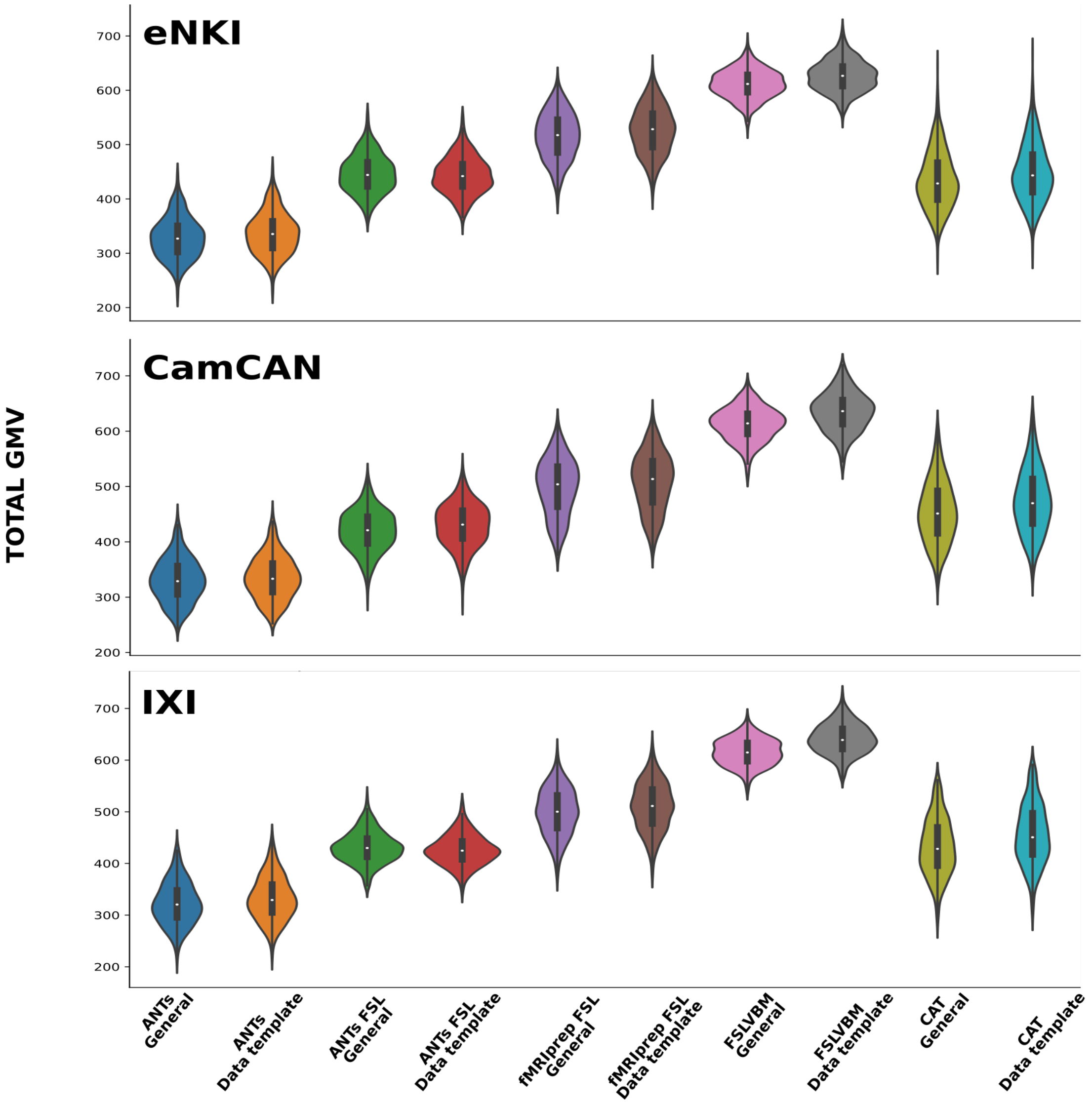


Figure 12: Total GMV of all subjects for all pipelines and all datasets. Important differences in the total intensities between all pipelines. Only ANTs-FSL and CAT have similar means. Not surprisingly, the template appears to have no impact on the total GMV of subjects. High consistency is observed for the same pipelines across datasets.

Classifying subjects' images based on the preprocessing pipeline. We used two methods to scale the features prior to classification, using a linear SVM: standard scaling and MinMax scaling. The classification results were close to perfect using both methods (Figure 13).

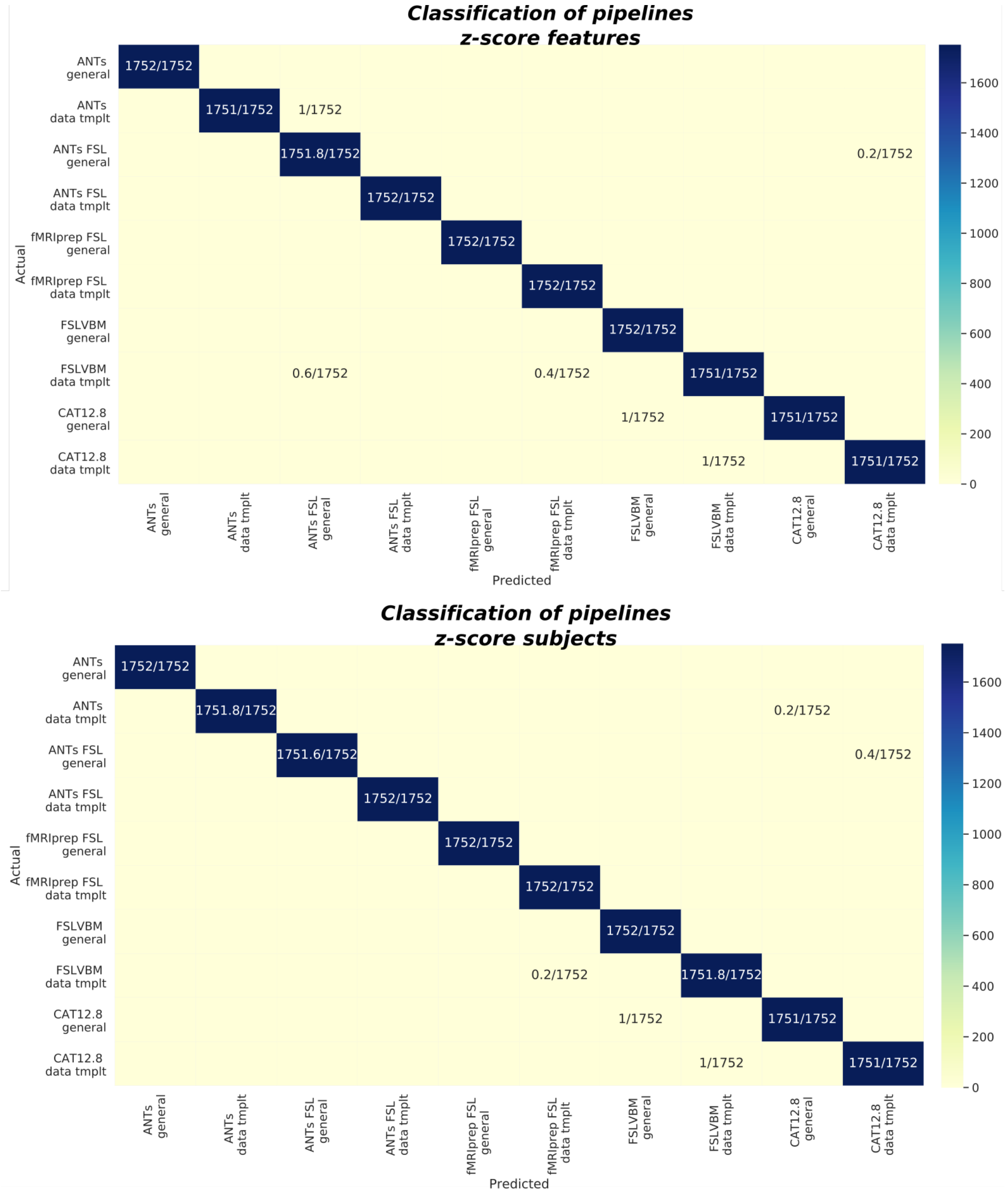


Figure 13: Confusion matrices of multiclass classification of preprocessed subjects from all pipelines using the preprocessing pipelines as labels. We tried two methods for scaling the features aiming at ruling out that the overall intensity differences drive the classification. We used standard scaling, which standardizes features by removing the mean and scaling to unit variance, and MinMax scaling, which scales and translates each feature individually such that it is in the given range on the training set, here between zero and one.

- **Similarity between pipelines** as expressed by the regionwise Pearson's correlation across subjects for each pair of pipelines. For CamCAN in figure 15 and for IXI in 16. From the figures of the three datasets, we see that similarities are consistent across datasets. However, some lack of variability is still identified, most likely due to differences in the quality of the images among datasets.

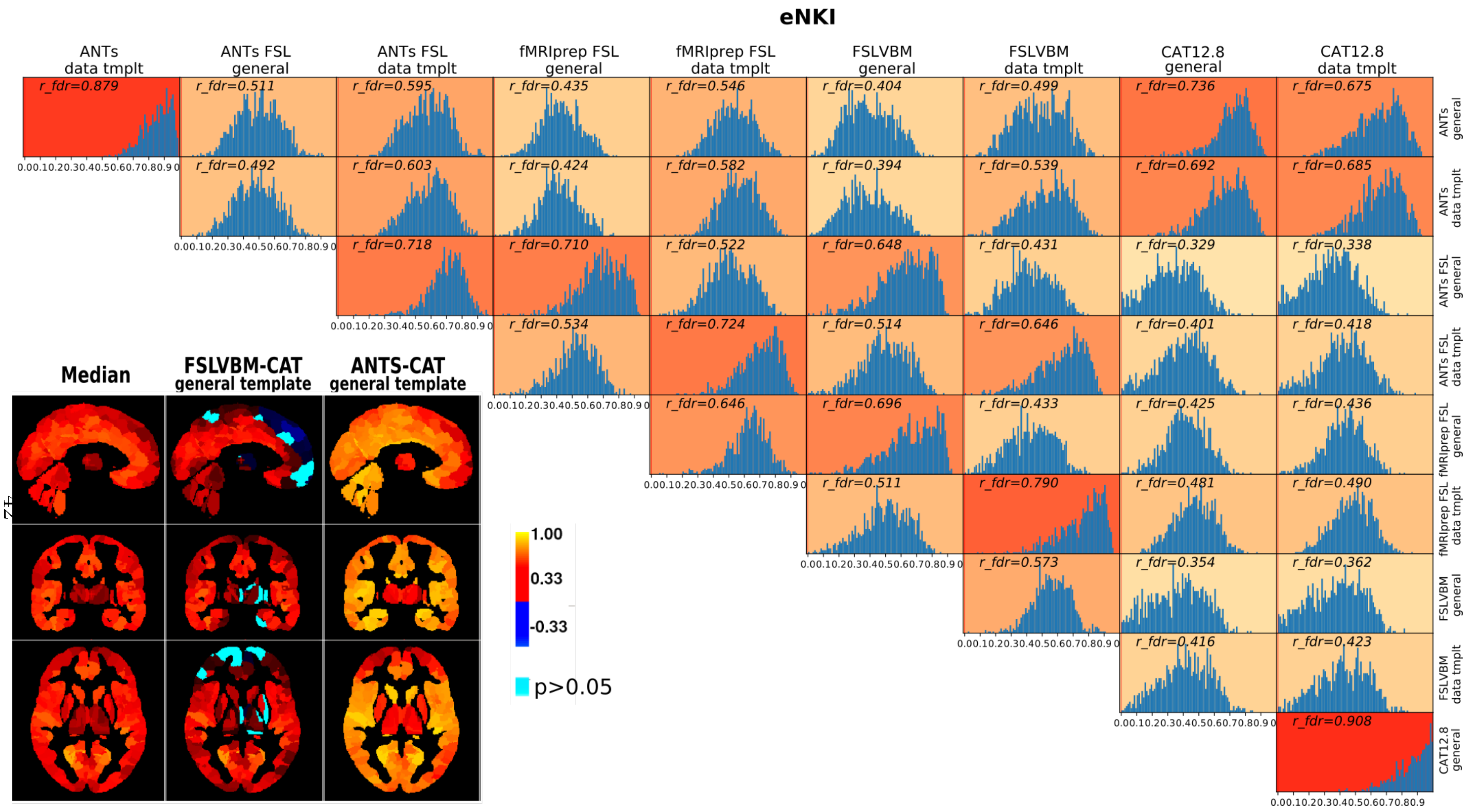


Figure 14: eNKI: Histograms of Pearson's R values for all regions across subjects and for all combinations of pipelines. Niftis represent R values in the brain for the pipelines with Max mean, Min mean and the comparison of the two pipelines with the highest correlation to age.

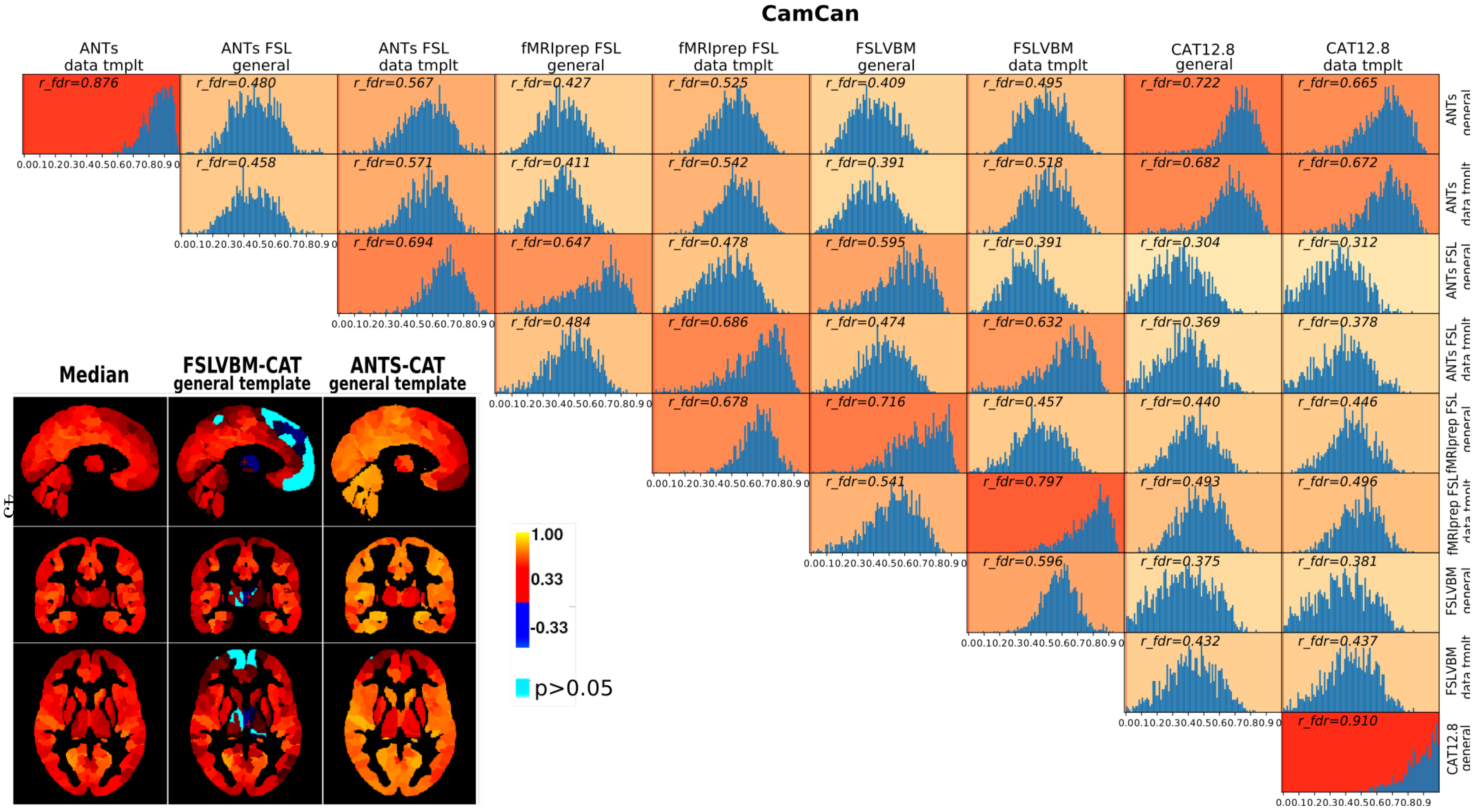


Figure 15: CamCAN: Histograms of Pearson's R values for all regions across subjects and for all combinations of pipelines. Niftis represent R values in the brain for the pipelines with Max mean, Min mean and the comparison of the two pipelines with the highest correlation to age.

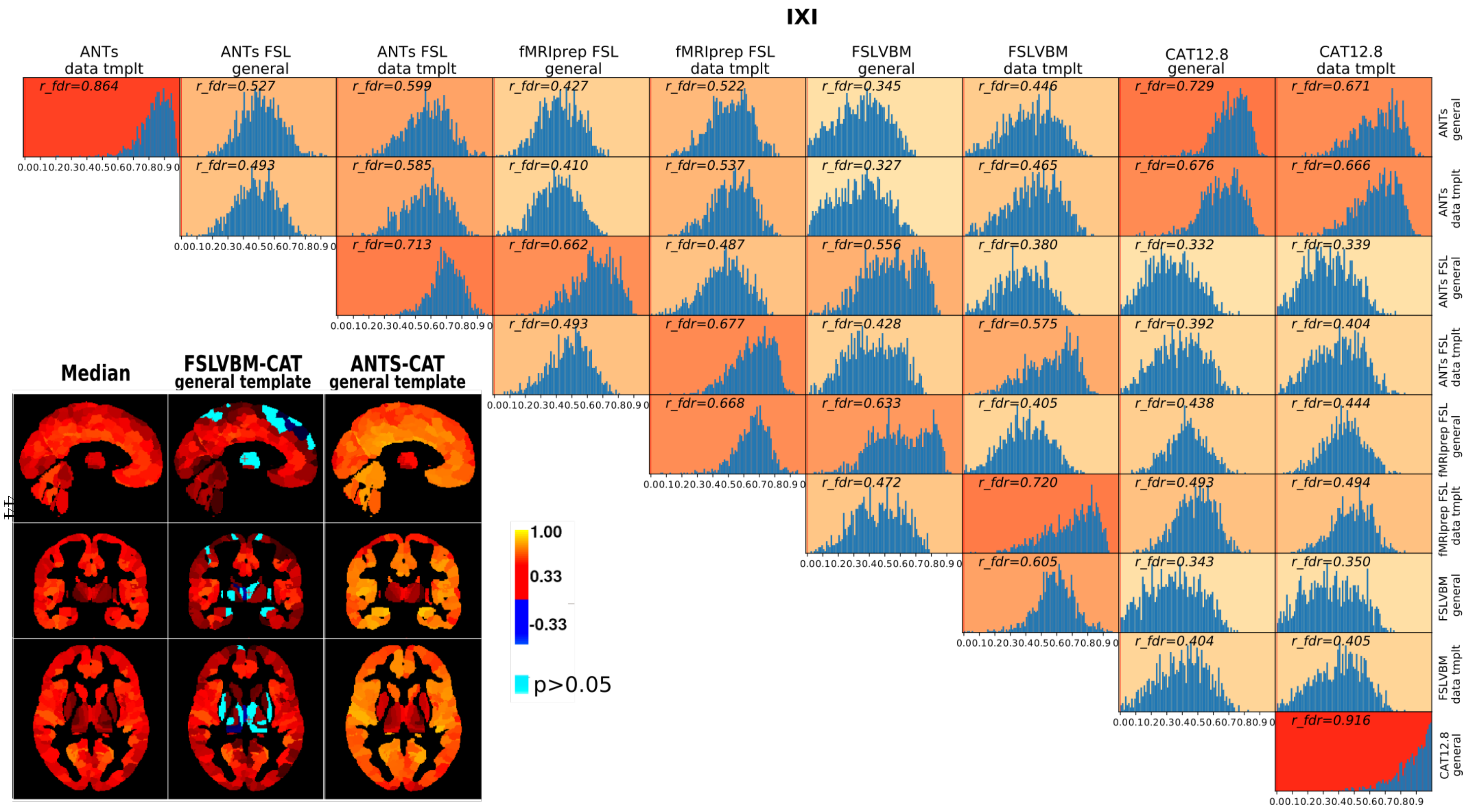




Figure 16: IXI: Histograms of Pearson's R values for all regions across subjects and for all combinations of pipelines. Niftis represent R values in the brain for the pipelines with Max mean, Min mean and the comparison of the two pipelines with the highest correlation to age.

Correlations of pipelines that differ only in the template for the three datasets 17

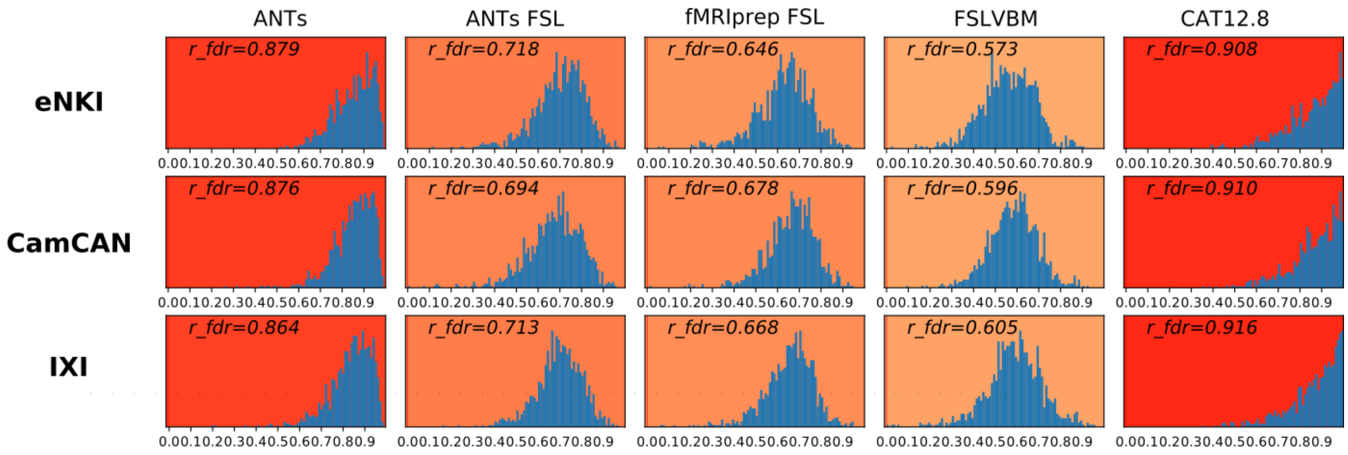


Figure 17: Mean correlation of regions across subjects for all datasets between pipelines that only differ in the template used for spatial normalization.

**The correlation between differential identifiability and Pearson's correlations** calculated between pairs of pipelines was examined to assess the agreement between the two methods. The results can be seen in Figure 18

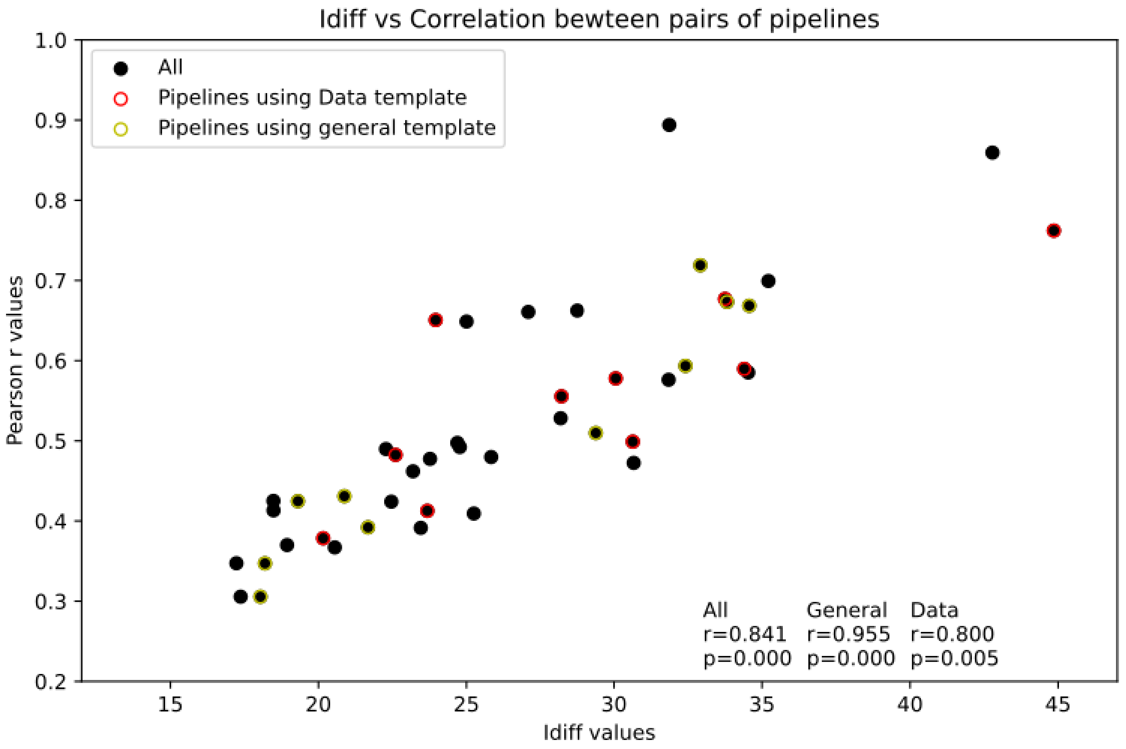


Figure 18: Strong correlations were observed between the two methods we used to assess similarity between pipelines, the univariate analysis and identification. Especially between the pipelines using the general template, the correlation was $r = 0.955$. For pipelines using data-templates, the correlation was $r = 0.8$. The correlation for all pipeline pairs was $r = 0.841$. All correlations had $p < 0.05$.

Age-ROIs correlations for all pipelines 19

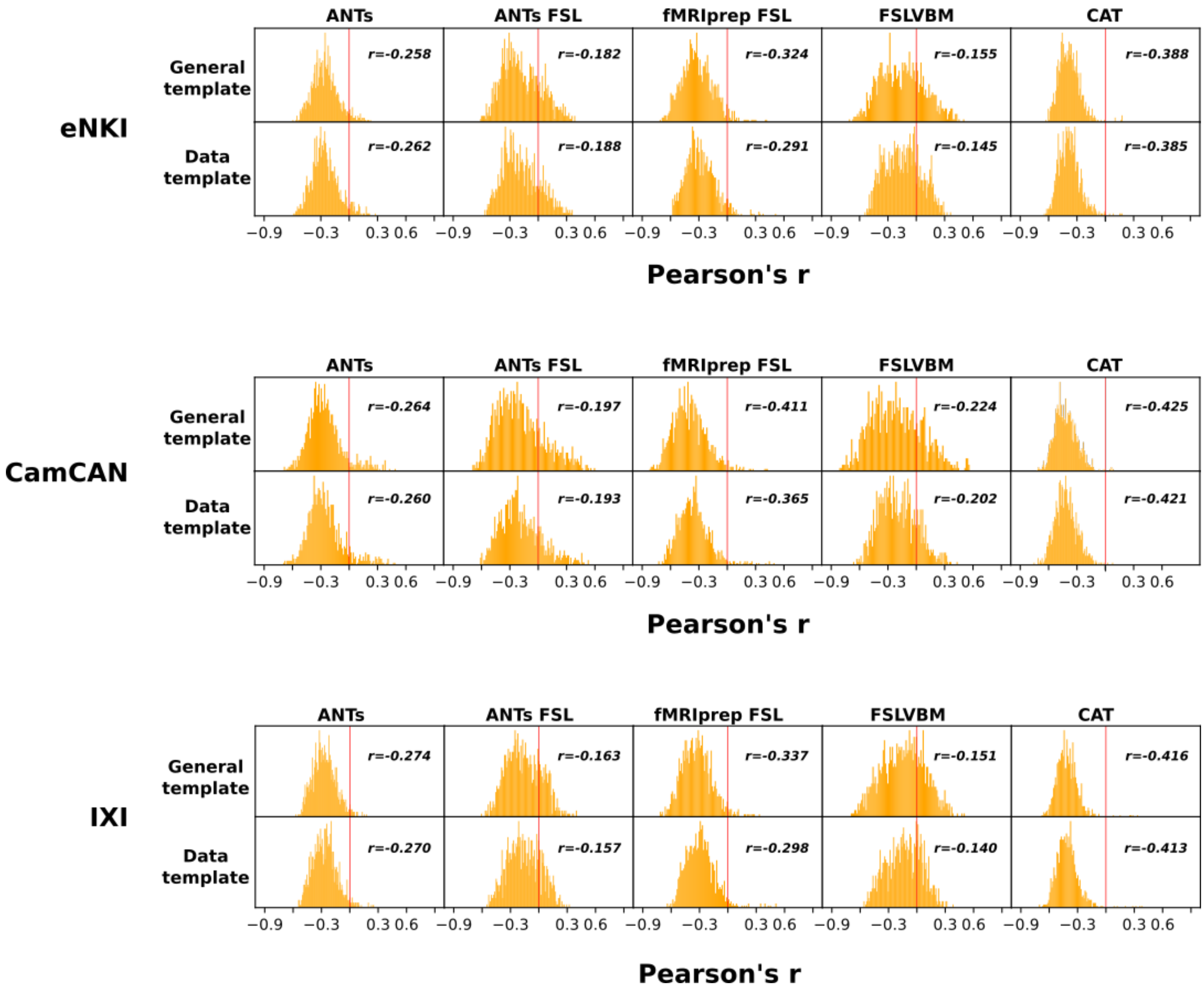


Figure 19: Correlation between regions and age of subjects for all pipelines.

Table 4 ANOVA results for correlation values between ROIs and age for all pipelines across subjects of all datasets. Figure 20 depicts the same correlation values between all regions and age calculated across subjects of all datasets, per pipeline.

| Dataset | F score | p-value |
|---|---|---|
| **eNKI** | 509.99 | 4.18E-193 <0.05 |
| **CamCAN** | 400.45 | 5.11E-156 <0.05 |
| **IXI** | 637.771 | 5.18E-234 <0.05 |
| **All data** | 324.468 | 5.18E-234 <0.05 |

Table 4: One-way ANOVA for the three datasets was performed for the three pipelines that used general templates and had the highest overall correlation to age.

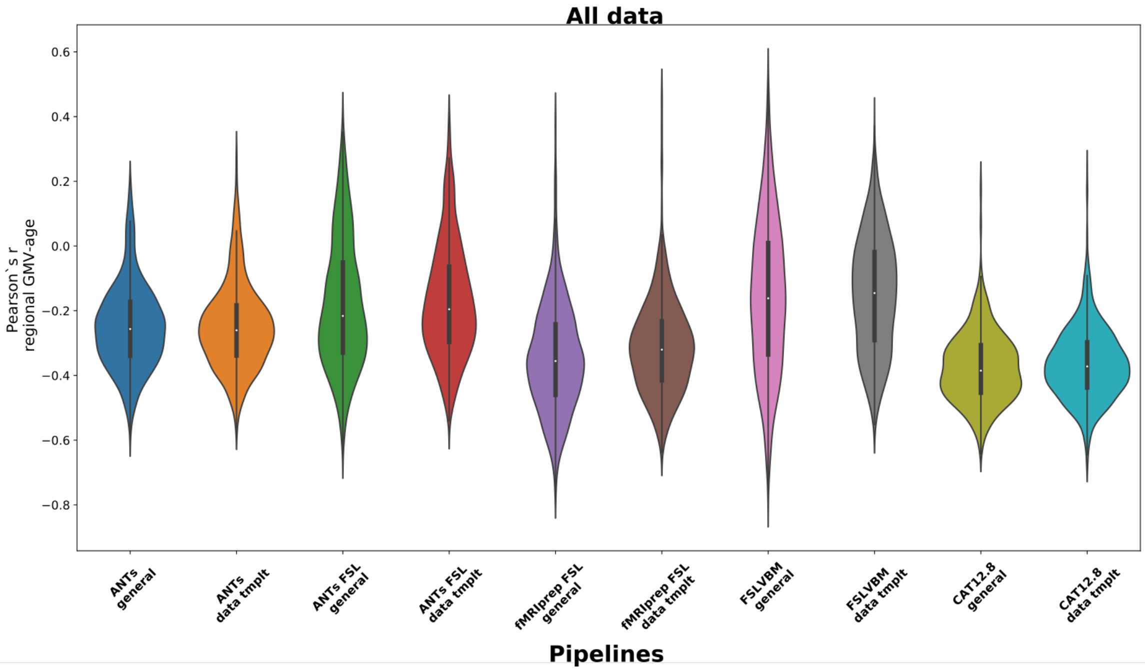


Figure 20: ROI-age correlations for all pipelines and all datasets. One-way ANOVA showed that there were significant differences between pipelines in ROI-Age correlations.

Scatter plots for each pipeline demonstrate the size of each ROI on the x-axis and the Pearson's r value between ROI and age calculated across subjects. The first figure (Figure 27) is for the CamCAN dataset, and Figure 28 is for IXI.

Paired comparisons of ROI-Age correlation values between pipelines

Figure 21 shows pairplots of age-region correlations across subjects for all data, and figures 22, 23 and 24 show for eNKI, CamCAM and IXI, respectively.

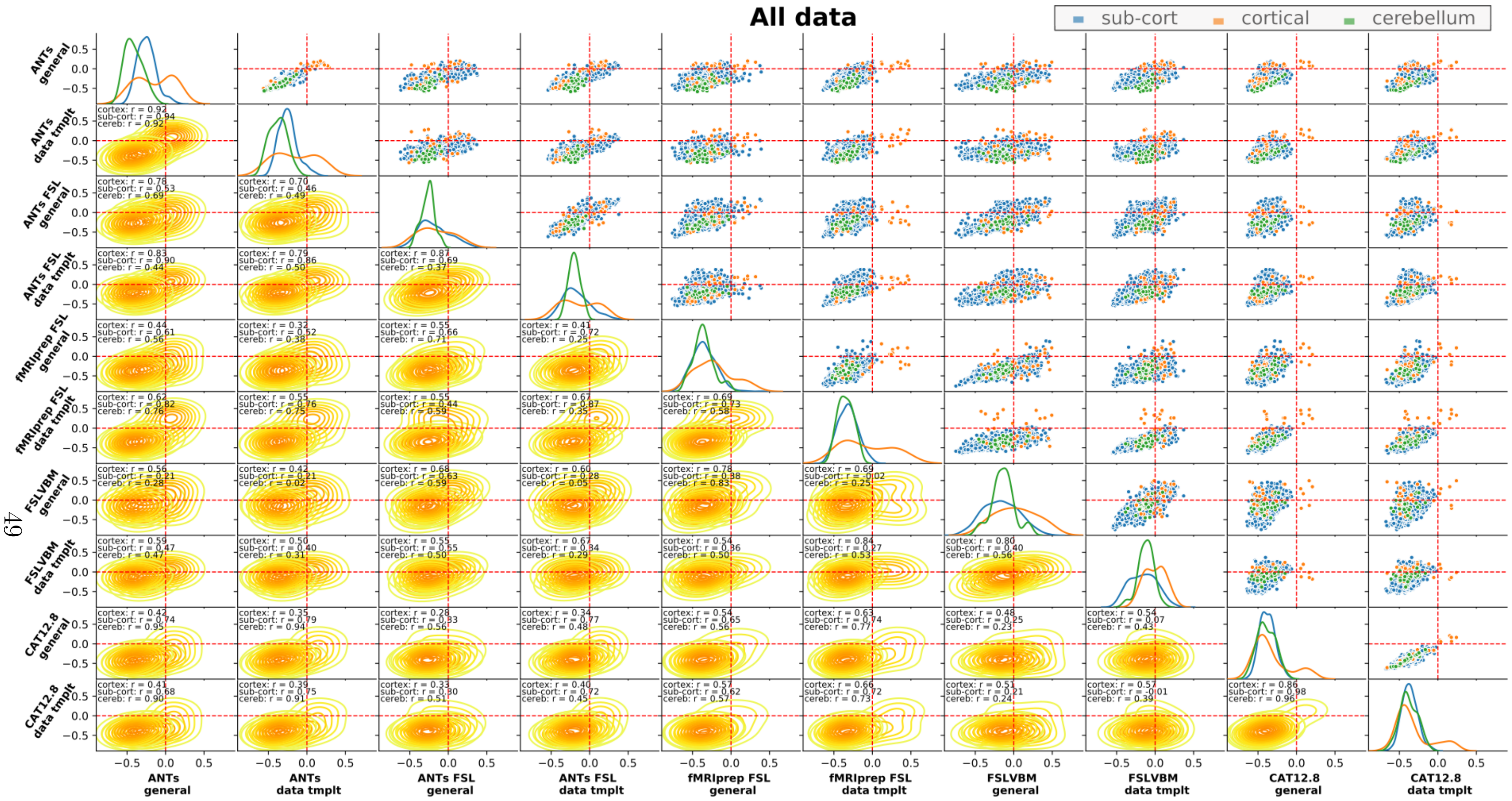
All data
sub-cort
cortical
cerebellum
ANTs general
ANTs data tmplt
ANTs FSL general
ANTs FSL data tmplt
fMRIprep FSL general
fMRIprep FSL data tmplt
FSLVBM general
FSLVBM data tmplt
CAT12.8 general
CAT12.8 data tmplt

Figure 21:

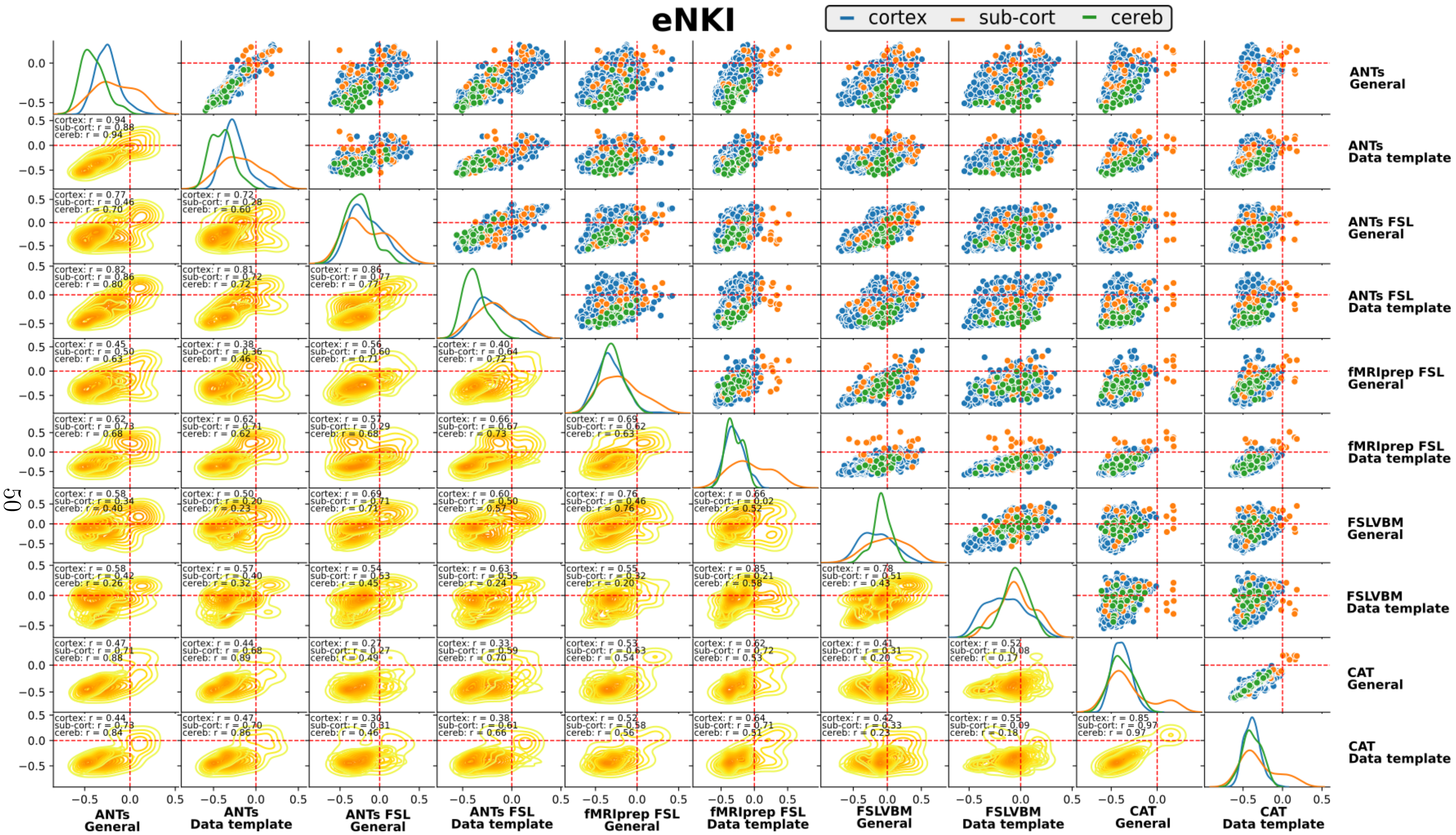
eNKI
cortex
sub-cort
cereb
ANTs General
ANTs Data template
ANTs FSL General
ANTs FSL Data template
fMRIprep FSL General
fMRIprep FSL Data template
FSLVBM General
FSLVBM Data template
CAT General
CAT Data template

Figure 22:

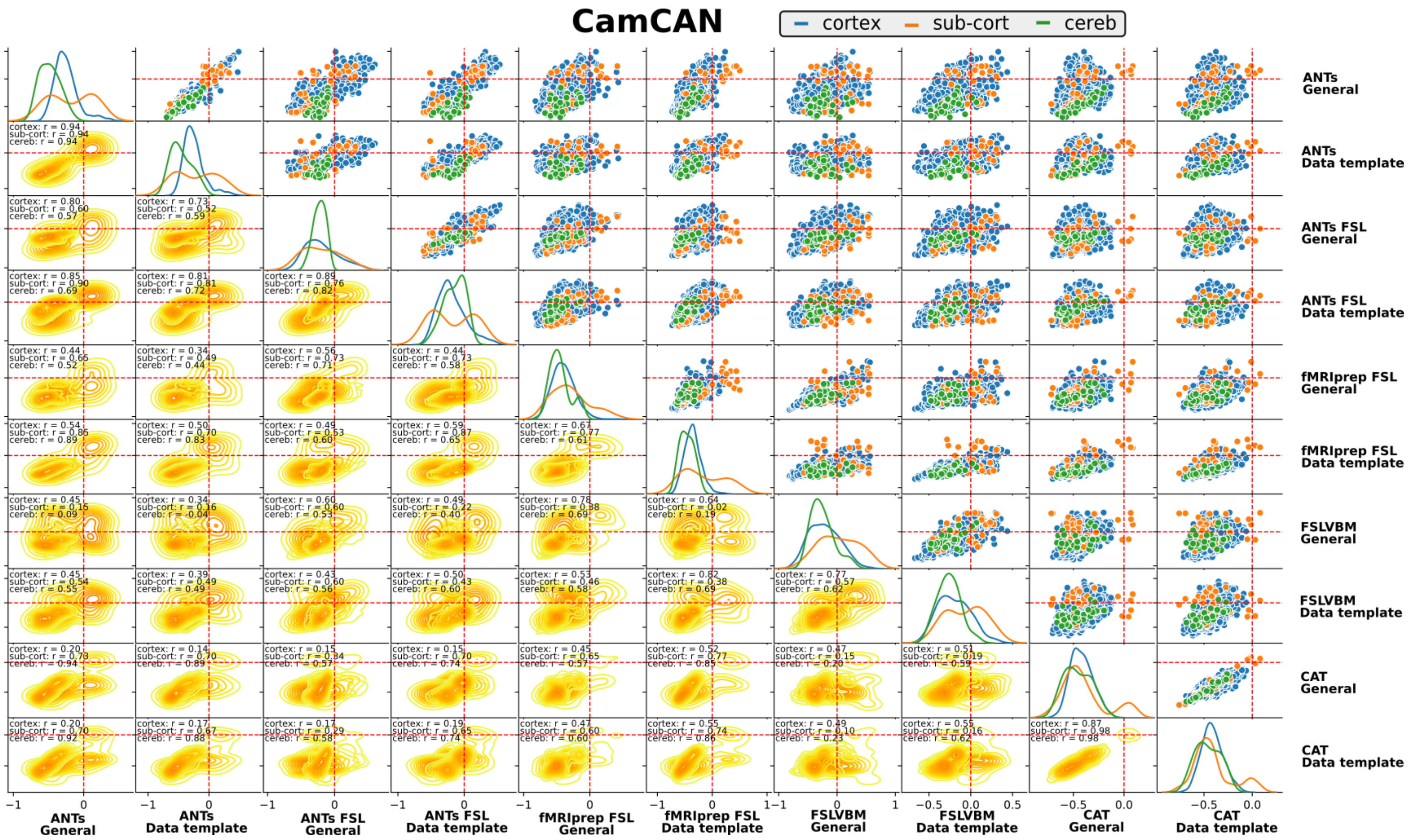
CamCAN
cortex
sub-cort
cereb
ANTs General
ANTs Data template
ANTs FSL General
ANTs FSL Data template
fMRIprep FSL General
fMRIprep FSL Data template
FSLVBM General
FSLVBM Data template
CAT General
CAT Data template

Figure 23:

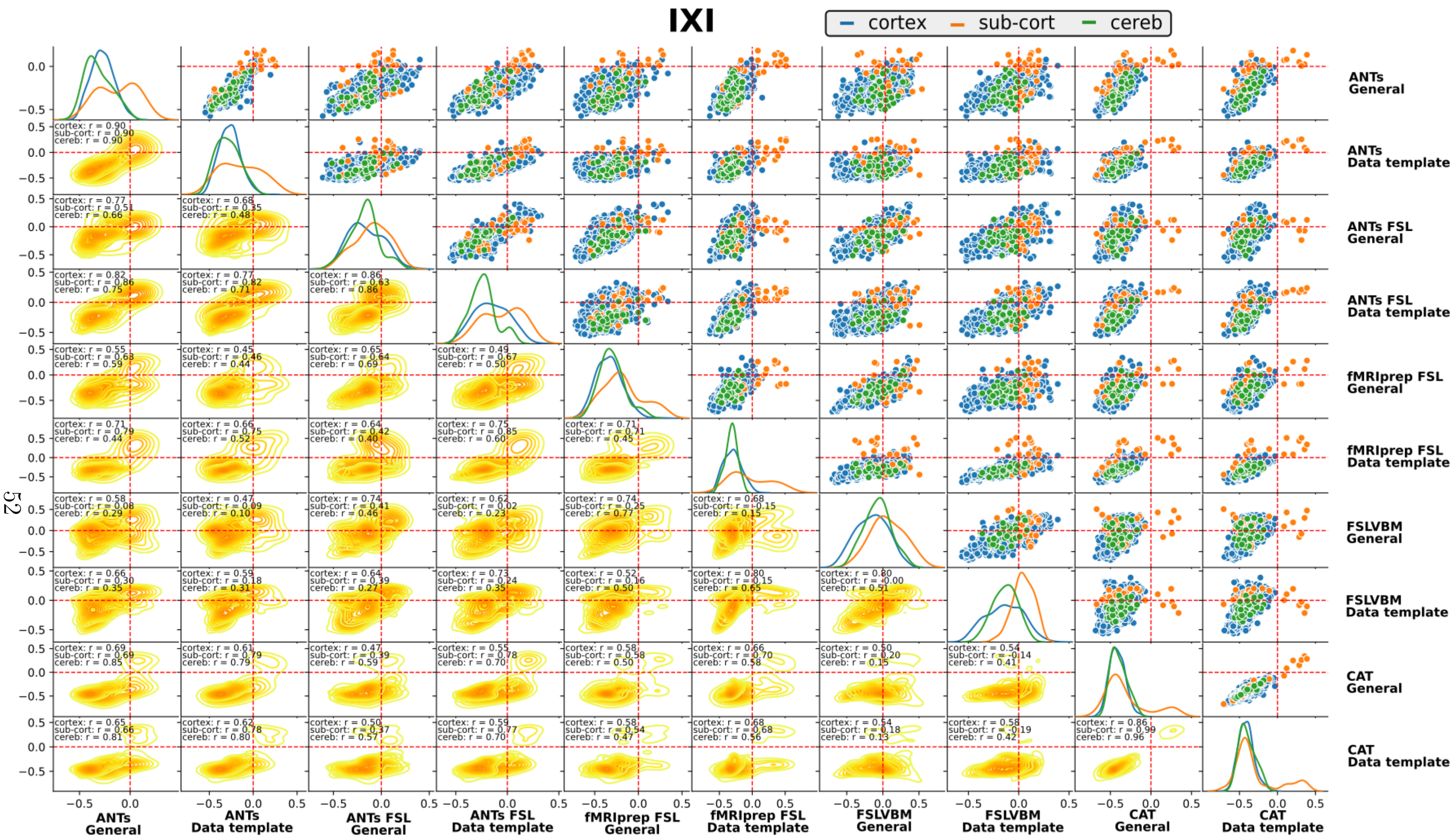
IXI
cortex
sub-cort
cereb
ANTs General
ANTs Data template
ANTs FSL General
ANTs FSL Data template
fMRIprep FSL General
fMRIprep FSL Data template
FSLVBM General
FSLVBM Data template
CAT General
CAT Data template

Figure 24:

The effect of region size

- Association between the overall similarity among the pipelines (calculated as the median of agreement between pairs) and parcel sizes

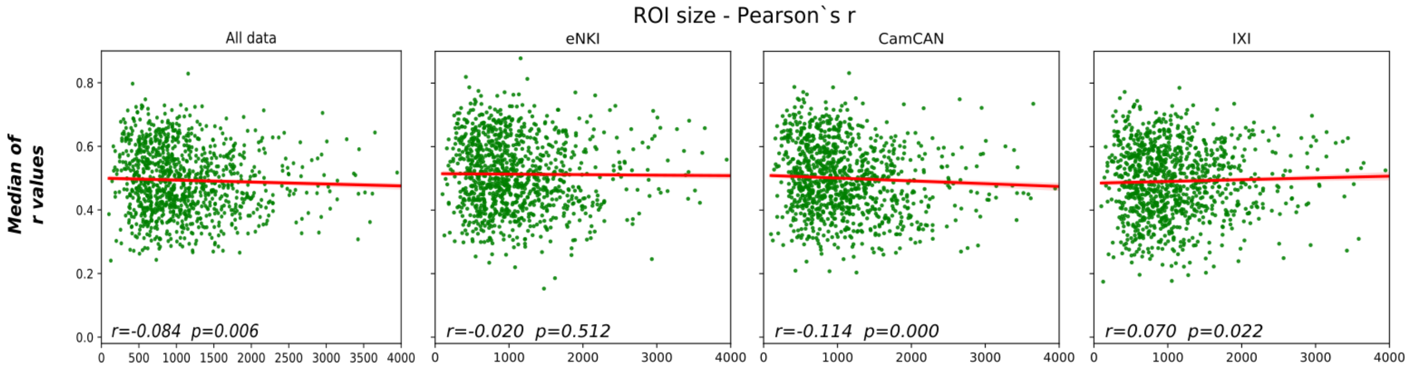


Figure 25: Median values from all pairs of pipelines of Pearson's r correlations across subjects for all regions plotted against the size of the regions. A nonsignificant correlation was found for the eNKI dataset, and very low correlations were found for the other two datasets.

-The association between the size of regions and the corresponding ROI-age correlation values. CAT appears to have a higher association between the size of each ROI and the correspondence correlation value with age for eNKI (r=-0.128 for both templates) and CamCAN. ANTs had similar values but only for the eNKI dataset (=r=-0.121 for general template and -0.125 for data template). For the IXI dataset, FSLVBM with the general template had the highest values (r=0.102). Notably, FSLVBM had a positive correlation when ANTs and CAT had negative values. Figure 29 provides the same analysis when data from all the datasets are combined.

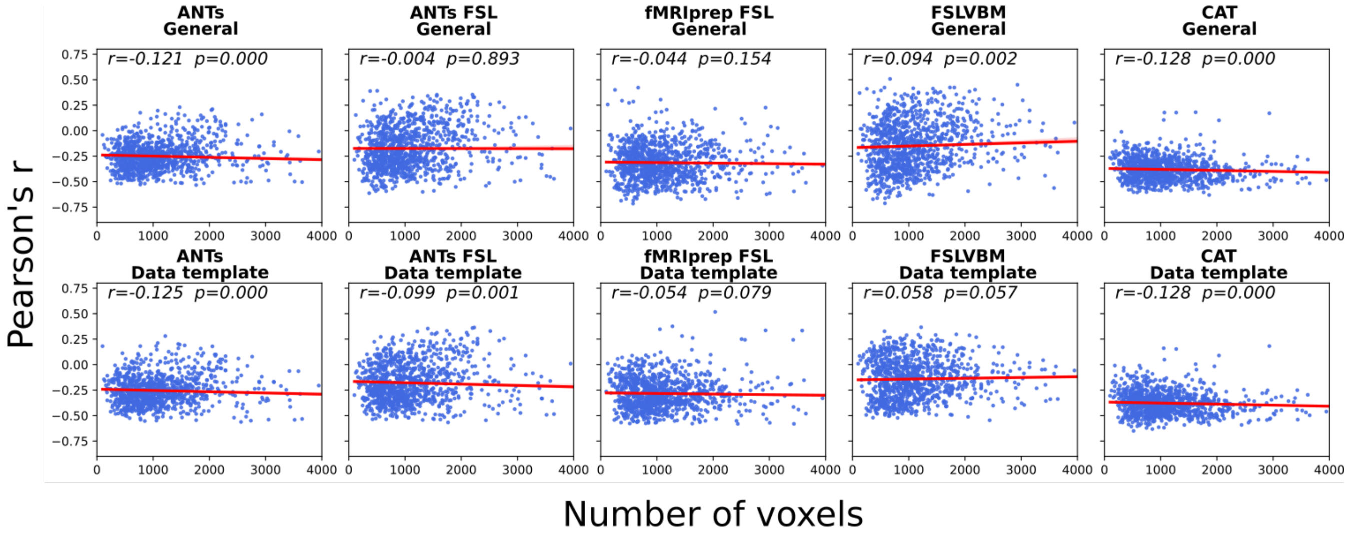


Figure 26: Scatter plots with the y-axis representing regional correlation to age and the x-axis representing the size of the corresponding ROI, for all pipelines estimated in all datasets. For each pipeline, we estimated the Pearson's r and p values. Red lines represent the linear regression line.

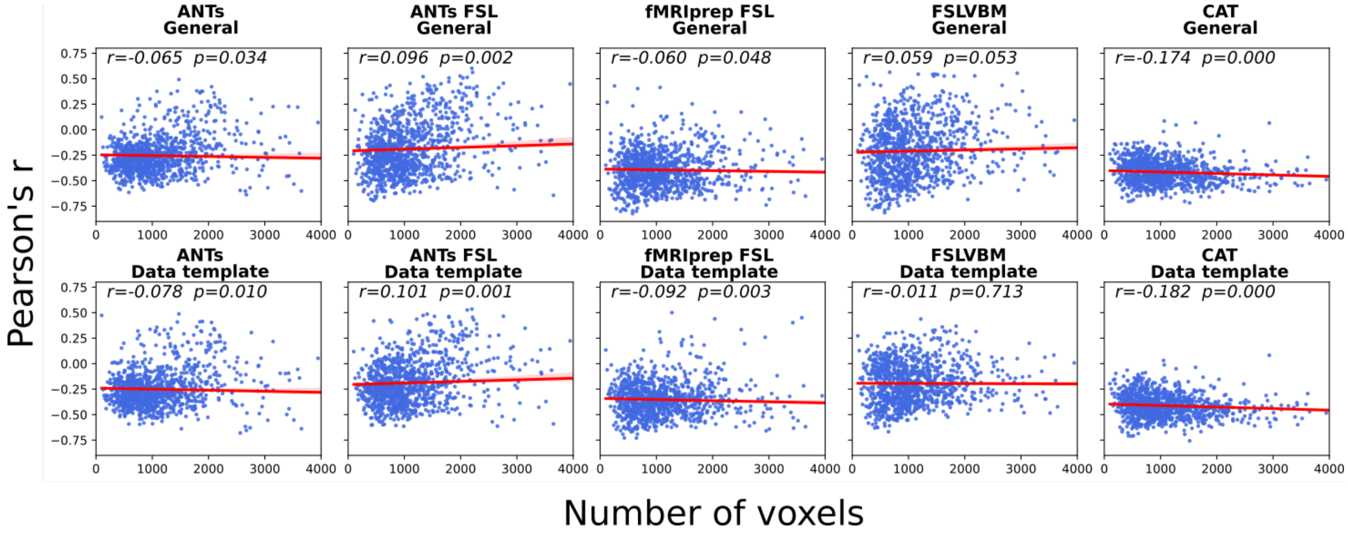


Figure 27: Scatter plots with the y-axis representing regional correlation to age and the x-axis representing the size of the corresponding ROI for all pipelines estimated in the eNKI dataset. For each pipeline, we estimated the Pearson's r and p values. Red lines represent the linear regression line.

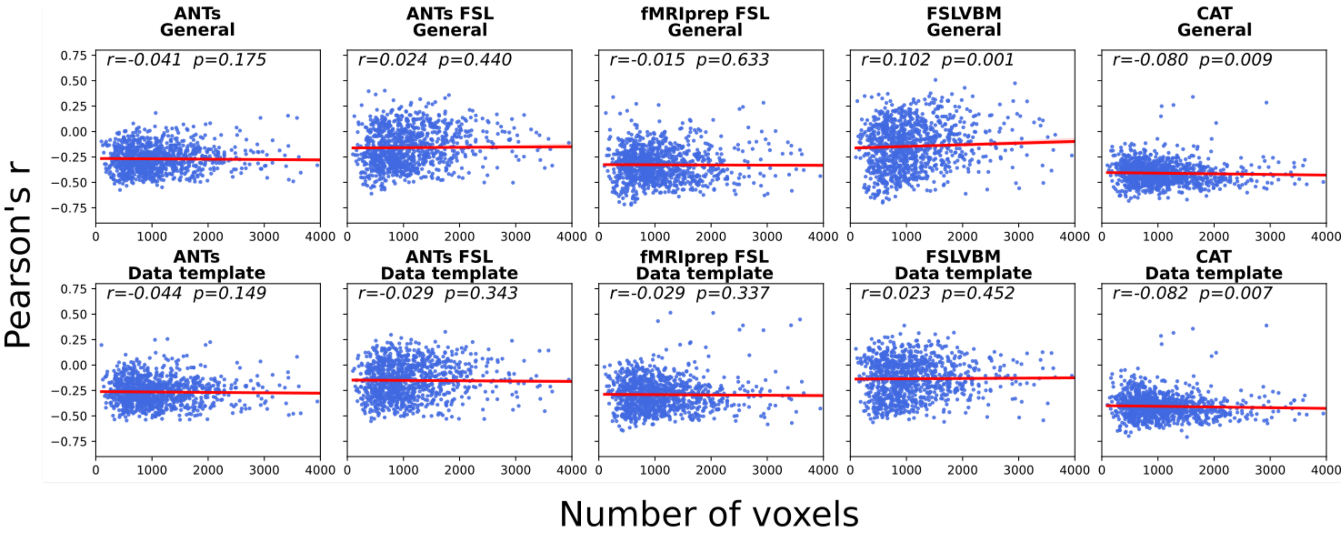


Figure 28: Scatter plots with the y-axis representing regional correlation to age and the x-axis representing the size of the corresponding ROI for all pipelines estimated in the CamCAN dataset. For each pipeline, we estimated the Pearson's r and p values. Red lines represent the linear regression line.

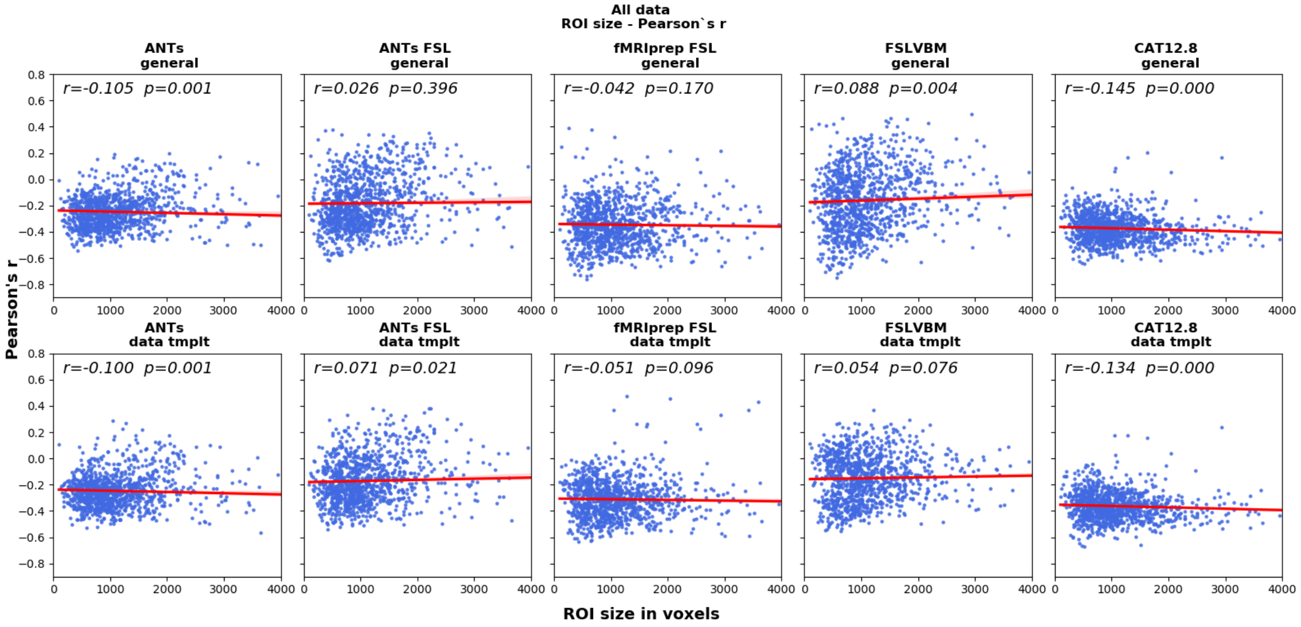


Figure 29: Scatter plots with the y-axis representing regional correlation to age and the x-axis representing the size of the corresponding ROI for all pipelines estimated in the IXI dataset. For each pipeline, we estimated the Pearson's r and p values. Red lines represent the linear regression line.

Detailed results of age predictions

Table 5 shows the analytical results of age prediction when models are trained and tested within each site.

Table 6 shows the analytical results of age prediction when the models are trained with two of the datasets and tested with the leftout dataset.

| **Pipelines** | **Models** | **MAE per model all datasets** | **MAE all models all datasets** |
|---|---|---|---|
| ANTs General template | RVR | 6.9 | **7.06** |
| | GPR | 7.04 | |
| | LASSO | 7.32 | |
| | KRR | 7.00 | |
| ANTs Data template | RVR | 6.93 | **7.04** |
| | GPR | 6.93 | |
| | LASSO | 7.35 | |
| | KRR | 6.95 | |
| ANTs FSLVBM General template | RVR | 6.56 | **6.55** |
| | GPR | 6.34 | |
| | LASSO | 6.79 | |
| | KRR | 6.52 | |
| ANTs FSLVBM Data template | RVR | 6.71 | **6.74** |
| | GPR | 6.46 | |
| | LASSO | 7.08 | |
| | KRR | 6.71 | |
| fMRIPrep FSL General template | RVR | 5.92 | **5.83** |
| | GPR | 5.65 | |
| | LASSO | 6.15 | |
| | KRR | 5.59 | |
| fMRIPrep FSL Data template | RVR | 6.14 | **6.18** |
| | GPR | 6.01 | |
| | LASSO | 6.51 | |
| | KRR | 6.06 | |
| FSLVBM General template | RVR | 6.25 | **6.17** |
| | GPR | 5.93 | |
| | LASSO | 6.45 | |
| | KRR | 6.05 | |
| FSLVBM Data template | RVR | 6.60 | **6.55** |
| | GPR | 6.30 | |
| | LASSO | 6.77 | |
| | KRR | 6.54 | |
| CAT 12 General template | RVR | 6.48 | **6.39** |
| | GPR | 6.26 | |
| | LASSO | 6.45 | |
| | KRR | 6.37 | |
| CAT 12 Data template | RVR | 6.46 | **6.37** |
| | GPR | 6.19 | |
| | LASSO | 6.47 | |
| | KRR | 6.37 | |

Table 5: Results of age prediction using a multivariate approach. Four models were tested in the three datasets in a nested K-fold scheme. The third column contains the averaged results of the three datasets per model. The last column shows the average of all datasets and all models for each pipeline.

| **Pipeline** | **Models** | **Test eNKI** | **Test CamCAN** | **Test IXI** | **Mean test (datasets)** | **Mean test (datasets & Pipelines)** |
|---|---|---|---|---|---|---|
| ANTs General template | RVR | 7.39 | 8.43 | 8.23 | 8.02 | 7.86 |
| | GPR | 7.19 | 7.94 | 8.27 | 7.80 | |
| | LASSO | 7.08 | 7.88 | 8.33 | 7.76 | |
| | KRR | 7.40 | 7.87 | 8.28 | 7.85 | |
| ANTs Data template | RVR | 7.48 | 8.26 | 7.88 | 7.87 | 7.86 |
| | GPR | 7.80 | 7.74 | 7.53 | 7.69 | |
| | LASSO | 7.73 | 6.88 | 8.99 | 7.87 | |
| | KRR | 7.84 | 8.12 | 8.03 | 8.00 | |
| ANTs-FSL General template | RVR | 6.96 | 9.16 | 9.75 | 8.62 | 8.44 |
| | GPR | 6.85 | 8.48 | 9.74 | 8.36 | |
| | LASSO | 7.60 | 7.97 | 9.88 | 8.49 | |
| | KRR | 6.73 | 7.91 | 10.28 | 8.31 | |
| ANTs-FSL Data template | RVR | 7.12 | 15.87 | 9.87 | 10.95 | 10.07 |
| | GPR | 6.95 | 11.47 | 9.72 | 9.38 | |
| | LASSO | 7.51 | 13.94 | 8.45 | 9.97 | |
| | KRR | 7.04 | 13.33 | 9.54 | 9.97 | |
| FMRIprep-FSL General template | RVR | 6.25 | 5.58 | 7.47 | 6.43 | 6.26 |
| | GPR | 6.10 | 5.63 | 7.13 | 6.29 | |
| | LASSO | 6.63 | 5.62 | 6.36 | 6.20 | |
| | KRR | 6.23 | 5.33 | 6.82 | 6.13 | |
| FMRIprep-FSL Data template | RVR | 6.83 | 6.56 | 6.12 | 6.50 | 6.21 |
| | GPR | 6.61 | 5.79 | 6.06 | 6.15 | |
| | LASSO | 6.66 | 5.76 | 5.91 | 6.11 | |
| | KRR | 6.48 | 5.89 | 5.82 | 6.06 | |
| FSLVBM General template | RVR | 6.62 | 6.02 | 8.66 | 7.10 | 6.77 |
| | GPR | 6.67 | 5.83 | 7.08 | 6.52 | |
| | LASSO | 7.23 | 6.09 | 6.88 | 6.73 | |
| | KRR | 6.43 | 5.77 | 8.00 | 6.73 | |
| FSLVBM Data template | RVR | 11.28 | 10.67 | 7.63 | 9.92 | 9.36 |
| | GPR | 11.42 | 10.66 | 6.70 | 9.55 | |
| | LASSO | 10.95 | 9.82 | 6.69 | 9.16 | |
| | KRR | 11.14 | 9.78 | 6.74 | 9.22 | |
| CAT12.8 General template | RVR | 6.75 | 6.53 | 6.39 | 6.55 | 6.45 |
| | GPR | 6.71 | 6.88 | 5.88 | 6.49 | |
| | LASSO | 6.75 | 6.62 | 5.92 | 6.43 | |
| | KRR | 6.55 | 6.58 | 5.82 | 6.32 | |
| CAT12.8 Data template | RVR | 6.83 | 6.81 | 6.25 | 6.63 | 6.76 |
| | GPR | 7.14 | 6.53 | 6.02 | 6.57 | |
| | LASSO | 7.20 | 7.51 | 7.52 | 7.41 | |
| | KRR | 7.01 | 6.06 | 6.19 | 6.42 | |

Table 6: Cross-dataset age prediction results. For each pipeline we trained four models using two of the datasets and predicted the age of the subjects on the third, left-out dataset. The optimal parameters for each model were selected using a 5-fold cross validation scheme in the training datasets. The second to last column contains the mean of each model across all combinations of the datasets for training and testing for each pipeline. The last column contains mean across models and dataset combinations for each pipeline.
